# Minimally disruptive optical control of protein tyrosine phosphatase 1B

**DOI:** 10.1101/776203

**Authors:** Akarawin Hongdusit, Peter H. Zwart, Banumathi Sankaran, Jerome M. Fox

## Abstract

Protein tyrosine phosphatases regulate a myriad of essential subcellular signaling events, yet they remain difficult to study in their native biophysical context. Here we develop a minimally disruptive optical approach to control protein tyrosine phosphatase 1B (PTP1B)—an important regulator of receptor tyrosine kinases and a therapeutic target for the treatment of diabetes, obesity, and cancer—and we use that approach to probe the intracellular function of this enzyme. Our conservative architecture for photocontrol, which consists of a protein-based light switch fused to an allosteric regulatory element, preserves the native structure, activity, and subcellular localization of PTP1B, affords changes in activity that match those elicited by post-translational modifications inside the cell, and permits experimental analyses of the molecular basis of optical modulation. Findings indicate, most strikingly, that small changes in the activity of PTP1B can cause large shifts in the phosphorylation states of its regulatory targets.

## INTRODUCTION

The enzymatic phosphorylation of tyrosine residues is centrally important to cellular function. It controls the location and timing of cellular differentiation, movement, proliferation, and death^1–4^; its misregulation can cause cancer, diabetes, and neurodegenerative diseases, among other disorders^5–7^. Methods to control the activity of phosphorylation-regulating enzymes without interfering with their native structure or cellular organization could, thus, enable detailed analyses of the mechanisms by which cells process essential chemical signals^8, 9^.

Optogenetic actuators—genetically encoded proteins that undergo light-induced changes in conformation—provide a powerful means of controlling enzyme activity over time and space. As protein fusion partners, they have enabled optical manipulation of biomolecular transport, binding, and catalysis with millisecond and submicron resolution^10, 11^. Common strategies to integrate optogenetic actuators into enzymes include (i) attachment near an active site, where they control substrate access^12, 13^, (ii) insertion within a catalytic domain, where they afford activity-modulating structural distortions^14^, and (iii) fusion to N- or C-termini, where they direct subcellular localization^15^ or guide domain assembly^16^. These approaches have generated powerful tools for stimulating phosphorylation-mediated signaling networks; unfortunately, their reliance on disruptive structural modifications has tended to limit their use in biochemical studies of native regulatory effects, for example, spatially dependent protein-protein interactions or, more notably, shifts in activity that match, rather than artificially exceed, those caused by post-translational modifications of an enzyme under study.

Protein tyrosine phosphatase 1B (PTP1B) is an important regulatory enzyme for which minimally disruptive architectures for photocontrol could prove particularly informative. This enzyme, which catalyzes the hydrolytic dephosphorylation of tyrosine residues, helps regulate insulin, leptin, and epidermal growth factor signaling and participates in a diverse set of spatiotemporally complex signaling processes^17^. PTP1B has two intriguing biophysical traits that make it particularly amenable to optogenetic study: (i) Its catalytically essential WPD loop undergoes cyclic, open-and-close motions that control the rate of phosphotyrosine hydrolysis at the active site^18^, and its C-terminal α7 helix modulates these motions through an allosteric network that extends over 25 Å across the protein (Fig. 1a). An architecture for photocontrol that makes use of this network could afford changes in activity that preserve interactions between PTP1B and its regulatory targets. (ii) PTP1B undergoes several post-translational modifications outside of its active site that cause modest, yet physiologically influential shifts in its activity (i.e., 1.7-3.1 fold^19, 20^; Supplementary Table 1). An optogenetic construct that affords similar changes in activity could help determine whether or not they—rather than the specific post-translational modifications that cause them—are sufficient to influence cellular physiology.

**Figure 1.**
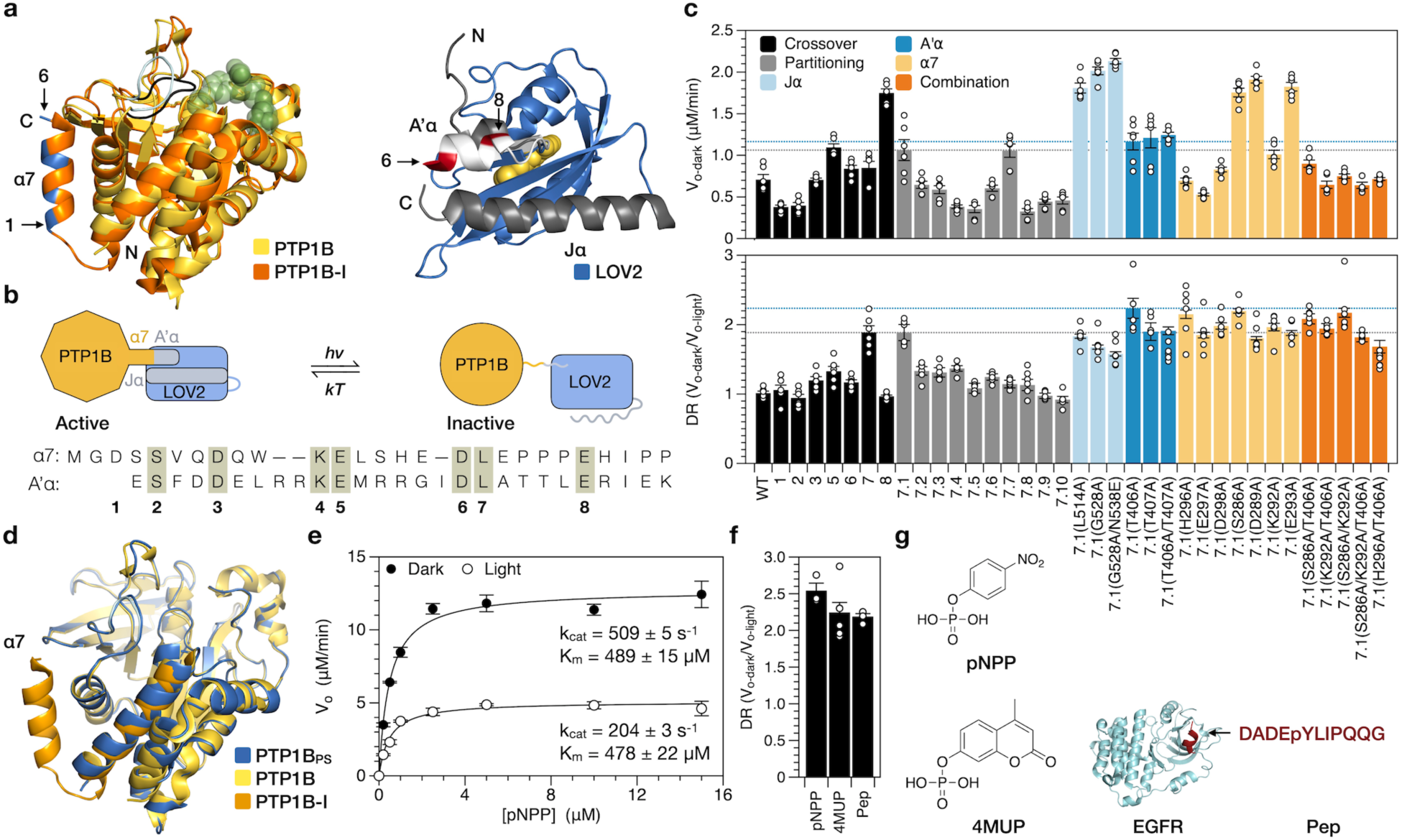
Minimally disruptive photocontrol of PTP1B. **a**, Left: An alignment of a competitively inhibited structure of PTP1B (orange, pdb entry 2f71) and an apo structure of PTP1B (yellow, pdb entry 3a5j) highlight an allosteric control system. Closure of the WPD loop (black) over an inhibitor orders the α7 helix; opening of the loop (red) hinders this ordering. Right: An alignment of the LOV2 domain from *A. sativa* (blue) and an N-terminal segment of the same domain of *A. thaliana* (white) that is identical between the two proteins (pdb entries 2v0w and 4hhd, respectively). Two terminal α-helices (gray and white) are stable in the dark state, but not the light state. **b**, Design of a photoswitchable chimera. Light-induced unwinding of the A’α helix of LOV2 destabilizes the α7 helix of PTP1B, causing an allosteric conformational change that inhibits catalysis. We attached the C-terminal α7 helix of PTP1B to the N-terminal A’α helix of LOV2 at crossover points in a primary sequence alignment (1-8). These points are highlighted in blue (PTP1B) and red (LOV2) in **a**. **c**, Assays on 4-methylumbelliferyl phosphate (4MUP) show the results of chimera optimization. Construct 7 has the largest dynamic range (DR) of the crossover variants; 7.1 has a higher activity than 7, and 7.1(T406A), termed PTP1B_PS_, has a larger DR than 7.1. The dashed gray and blue lines denote values for 7.1 and 7.1(T406A), respectively. The plotted data depict the mean, SE, and associated estimates of DR for n = 6 independent experiments. **d,** Aligned catalytic domains of PTP1B in three structures: photoswitchable (6ntp), apo (3a5j), and competitively inhibited (2f71, α6 and α7 only). **e**, An analysis of the activity of PTP1B_PS_ on p-nitrophenyl-phosphate (pNPP) indicates that light affects *k_cat_*, but not *K_m_* (*k_cat-dark_*/*k_cat-light_* = 2.50 +/- 0.04). Error bars denote SE for n = 6 independent reactions. **f**, The DR of PTP1B_PS_ is similar for substrates of different sizes. The plotted data depict the mean, SE, and associated estimates of DR for n ≥ 3 independent reactions. **g**, Structures of pNPP, 4MUP, and a peptide (PEP) derived from epidermal growth factor receptor (EGFR). Source data are provided as a Source Data file.

In this study, we used a protein-based light switch to place the native allosteric regulatory system of PTP1B under optical control. This conservative optogenetic design preserved the native structure and subcellular localization of PTP1B, permitted changes in activity that match those caused by post-translational modifications inside the cell and, when paired with a FRET-based biosensor, enabled spatiotemporal control and measurement of intracellular PTP1B activity. An optogenetic analysis carried out with this system showed that small, transient changes in PTP1B activity can cause large shifts in insulin receptor (IR) phosphorylation. The optogenetic tools developed in this study thus complement existing methods for studying protein tyrosine kinases (PTKs)—which, unlike PTPs, possess many light-sensitive analogues and FRET-based biosensors^21, 22^—and, more broadly, demonstrate an optogenetic approach for studying regulatory enzymes in their native biophysical context.

## RESULTS

### Allosteric Photocontrol of PTP1B

We sought to place PTP1B under optical control by using LOV2, the light-sensitive domain from phototropin 1 of *Avena sativa*, to toggle the conformation of its α7 helix. LOV2 derives its optical activity from a noncovalently bound flavin mononucleotide (FMN) that, when exposed to blue light, forms an intermolecular carbon-sulfur bond that destabilizes the N- and C-terminal helices of the protein (Fig. 1a)^23^. We hypothesized that attachment of the N-terminal A’α helix of LOV2 to the C-terminal α7 helix of PTP1B would couple (i) light-induced unwinding of the A’α helix to (ii) destabilization of the α7 helix and disruption of WPD loop motions (Fig. 1b). To our satisfaction, several PTP1B-LOV2 chimeras—each generated by fusing the A’α and α7 helices at a different crossover point in a primary sequence alignment—exhibited light-dependent catalytic activity on 4-methylumbelliferyl phosphate (4MUP; Fig. 1c). Fusion of the Jα helix of LOV2 to the N-terminus of PTP1B, by contrast, did not confer photosensitivity (Supplementary Figures 1a-1b), a result consistent with the large distance between its N-terminus and active site (Fig. 1a).

To enhance the dynamic range (DR = V_o-dark_ / V_o-light_) of our most light-sensitive chimera (i.e., construct 7, where DR = 1.9), we used two approaches: First, we attempted to improve communication between the LOV2 and PTP1B domains by reducing the length of the linker between them; similar changes have improved photoswitching in other light-sensitive fusions^24^. Unfortunately, shorter linkers (which, through residue deletions, could also alter the phases of the fused helices) tended to reduce DR; we chose one variant with an unaltered DR—chimera 7.1—for further optimization. Next, we attempted to increase the stability of the dark state over the light state by adding stabilizing mutations to flexible helices. (For the Jα and A’α helices, we used established stabilizing mutaitons^23, 25^; for α7, we replaced solvent-exposed residues with alanine, which has a high helix propensity^26^). Stabilizing mutations in the Jα helix have amplified DRs of previous LOV2 fusions^25, 27^. Intriguingly, for our chimeras, mutations in the Jα helix improved activity, but reduced photosensitivity; several mutations in the A’α and α7 helices, by contrast, increased the DR (Fig. 1c). Overall, the effects of amino acid substitutions in these two helices were non-additive and reached a maximum DR of 2.2 on 4MUP. We chose a single high-DR chimera—7.1(T406A), termed PTP1B_PS_—for further study.

We assessed the structural integrity of the PTP1B domain within PTP1B_PS_ by using X-ray crystallography to examine its dark-state conformation. Intriguingly, although crystals of PTP1B_PS_ were yellow and turned clear when exposed to blue light—a behavior indicative of the presence of LOV2^12, 28^—diffraction data permitted placement and refinement of only PTP1B (Supplementary Figures 2 and 3). Detection of LOV2 was likely impeded by two interrelated crystallographic features: (i) a disordered α7 helix, which is unresolvable in apo structures of PTP1B^29^, and (ii) variability in the orientation of LOV2 within the crystal lattice (Supplementary Note 1). Despite this structural disorder, aligned catalytic domains of PTP1B_PS_ and wild-type PTP1B had a root-mean-square deviation of 0.30 Å (Fig. 1d). Crystallographic results, thus, suggest that LOV2 does not alter the native conformation of the catalytic domain of PTP1B.

We explored the mechanism of photomodulation by using kinetic assays to examine the influence of LOV2 on PTP1B-mediated catalysis. In brief, we measured the activity of PTP1B_PS_ on p-nitrophenyl-phosphate (pNPP) in the presence and absence of blue light (455 nm), and we used the initial rates to construct dark- and light-state Michaelis-Menten curves (Fig. 1e). These curves indicate that blue light reduces *k_cat_* by 2.5-fold but leaves *K_m_* unaltered. Data collected under repeated illumination, in turn, shows that changes in *k_cat_* are reversible (Supplementary Figure 4c). The isolated influence of LOV2 on *k_cat_* indicates that this photoswitch does not interfere with substrate binding and, importantly, is consistent with a mechanism in which LOV2 allosterically modulates WPD loop motions, which control the rate of hydrolysis^30^.

To assess the maximum achievable DR for our system, we removed the α7 helix of PTP1B; that is, we used an α7-less variant as a model for a maximally photoswitched form of the enzyme. Intriguingly, helix removal lowered *k_cat_* by 2.9-fold, suggesting that PTP1B_PS_ has a DR that is 85% of the maximum value for a photoswitch that inhibits catalytic activity by unwinding the α7 helix (Supplementary Figures 4a-4b). Importantly, this DR is within the range of DRs of previously developed light-sensitive signaling enzymes used to elicit physiologically relevant cellular responses to optical stimuli (DRs ∼1.7-10^12, 31^; Supplementary Table 8) and matches physiologically influential changes in activity caused by post-translational modifications of PTP1B that occur outside of its active site (i.e., phosphorylation, proteolysis, and sumoylation, which reduce/enhance PTP1B activity by 1.7-3.1 fold^19, 20^; Supplementary Table 1).

We hypothesized that an optogenetic system that exerts allosteric control over catalytically essential loop motions might—in contrast with a system that competitively inhibits the active site—exhibit a modulatory effect that is independent of substrate size and binding affinity. To test the substrate dependence of PTP1B_PS_, we measured its DR on pNPP, 4MUP, and a phosphorylated peptide (Fig. 1f). The DRs for these substrates differed by less than 15% (Fig. 1g); this similarity suggests that the magnitude of photocontrol is, in fact, substrate independent.

### Biophysical Analysis of Photocontrol

Although we designed our chimeras to exploit conformational changes in the N-terminal A’α helix of LOV2, the results of crystallographic and spectroscopic analyses of this photoswitch indicate that its N- and C-terminal helices work together to transduce conformational changes across the protein^23, 32^. To examine the contribution of both helices to the photoresponse of chimera 7.1, one of our most light-sensitive chimeras, we introduced disruptive mutations (i.e., we added charged residues at buried sites^32, 33^). For both helices, disruptive mutations reduced light-dependent catalytic activity as effectively as C450M, a “dark state” mutation that prevents the formation of the cysteine adduct in LOV2; complete removal of the Jα helix had the same effect (Fig. 2a). Our results thus indicate that both A’α and Jα helices are necessary for LOV2-mediated control of PTP1B activity.

**Figure 2.**
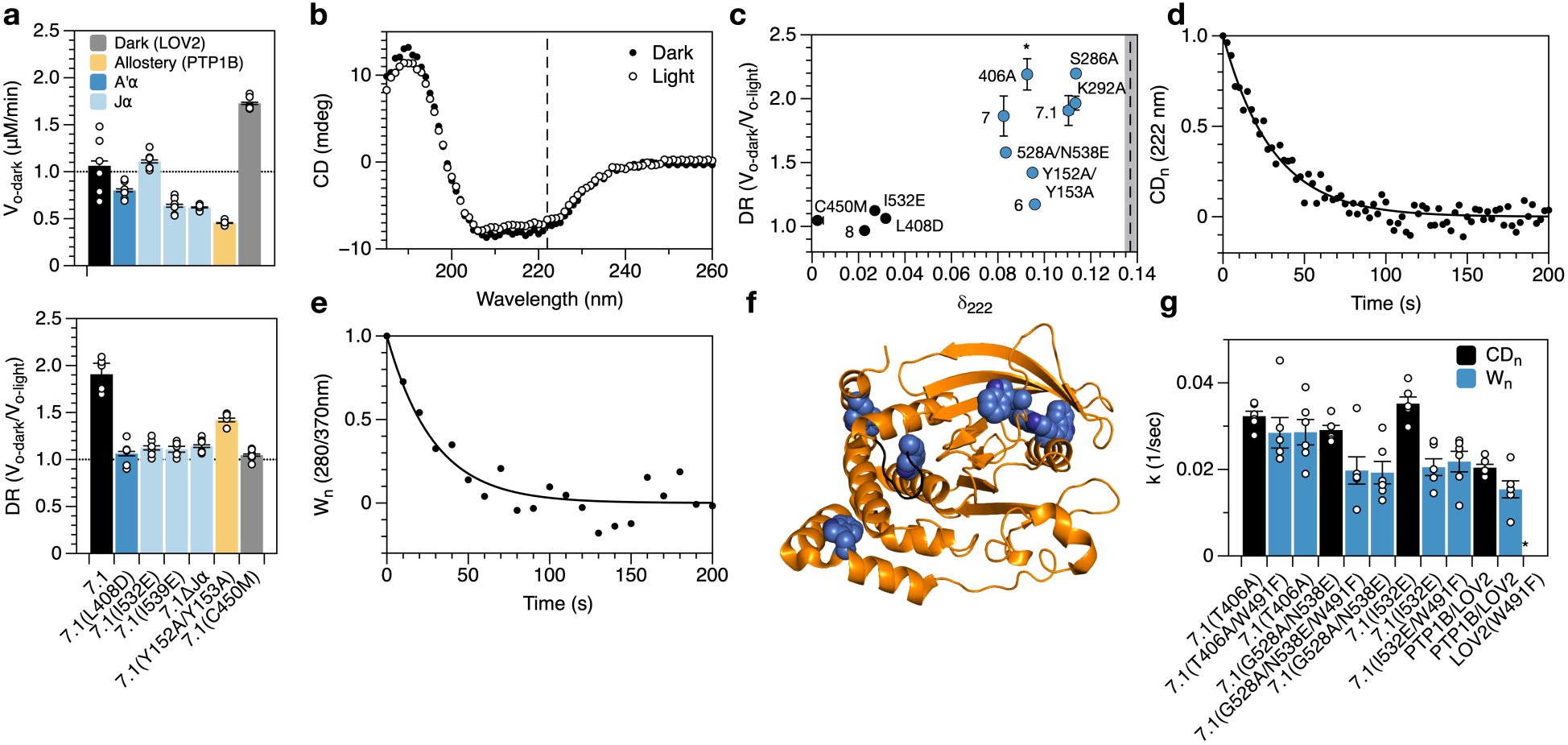
Analysis of allosteric communication in PTP1B_PS_. **a**, Mutations that either prevent adduct formation in LOV2 (C450M), destabilize the A’α and Jα helices (I532E, I539E, and ΔJα), or disrupt the allosteric network of PTP1B (Y152A/Y153A) reduce the photosensitivity of 7.1 and, with the exception of I532E and C450M, lower its specific activity. The plotted data depict the mean, SE, and associated estimates of DR for n = 6 independent experiments. **b**, Exposure of PTP1B_PS_ to 455 nm light reduces its α-helical content (i.e., the mean residue ellipticity [MRE] at 222 nm). **c**, An analysis of different chimeras indicates that light-induced changes in α-helical content (i.e., δ_222_ = [CD_222-dark_-CD_222-light_]/CD_222-dark_, or the fractional change in MRE at 222 nm) are necessary, but not sufficient for light-sensitive catalytic activity (i.e., high DR). Mutations correspond to variants of 7.1. Chimeras with large values of δ_222_ appear in blue; the dashed line indicates δ_222_ for equimolar amounts of free PTP1B and LOV2. Error bars denote SE for n = 6 independent reactions. **d-e**, Thermal recovery of (**d**) α-helical content (i.e., the change in MRE at 222 nm normalized by the full change over 250 seconds) and (**e**) tryptophan fluorescence (i.e., the change in fluorescence normalized by the full change over 250 seconds) of PTP1B_PS_. **f**, A crystal structure of PTP1B (pdb entry 2f71) shows the locations of six tryptophan residues (blue) and the WPD loop (yellow). **g**, Kinetic constants for thermal recovery are larger for α-helical content than for tryptophan fluorescence (the latter of which is not affected by W491). The discrepancy between these constants is smallest for PTP1B_PS_ (i.e., 7.1(T406A)). The plotted data depict the mean, associated data points, and SE for n = 6 independent reactions. Source data are provided as a Source Data file.

A previous NMR study of PTP1B dynamics showed that mutations in its L11 loop can disrupt allosteric communication between the α7 helix and WPD loop^29^. To confirm the contribution of allostery to photocontrol, we modified chimera 7.1 with a mutation known to exert such an effect: Y152A/Y153A. This modification reduced DR by ∼25%, a disruption distinct from the conservative/beneficial effects of alanine substitutions in the α7 helix (Fig. 2a). The sensitivity of DR to mutations in the L11 loop indicates that the native allosteric network of PTP1B is, indeed, necessary for optogenetic control of its catalytic activity.

We hypothesized that our most photoswitchable chimeras might exhibit large changes in secondary structure between light and dark states—changes that result largely from ordered-to-disordered transitions of the A’α, Jα, and α7 helices. To test this hypothesis, we used circular dichroism (CD) spectroscopy to compare optically induced shifts in α-helical content (δ222 = [CD222-dark-CD222-light]/CD222-dark; Fig. 2b). Intriguingly, changes were large for the chimeras with light-dependent catalytic activities but spanned a range of values for low-DR constructs, (Fig. 2c). The one-way dependence of DR on δ222 indicates that changes in α-helical conformation are necessary, but not sufficient for photocontrol.

We speculated that chimeras with large changes in α-helical content (δ_222_) but light-insensitive catalytic activities (low DRs) might suffer from weak conformational coupling between the LOV2 and PTP1B domains. To study this coupling, we carried out two experiments. In the first, we examined the thermal recovery of LOV2 from the light state by illuminating PTP1B-LOV2 chimeras with blue light and, subsequently, measuring the return of α-helical content in the dark (Fig. 2d). A link between the conformation of LOV2 and α-helical content is supported by (i) the sensitivity of δ_222_ to disruptive mutations in LOV2 and (ii) the insensitivity of δ_222_ to the catalytic response of PTP1B (that is, activity-modulating structural changes in PTP1B, which presumably differ between high- and low-DR chimeras, do not affect δ_222_). In the second experiment, we examined the thermal recovery of PTP1B by measuring the return of tryptophan fluorescence in the dark (Fig. 2e). A link between the conformation of PTP1B and tryptophan fluorescence is supported by (i) the existence of six tryptophan residues in PTP1B (Fig. 2f) and (ii) the insensitivity of recovery kinetics to the removal of W491, the only tryptophan in LOV2 (Fig. 2g). Intriguingly, kinetic constants for thermal recovery were higher for α-helical content than for tryptophan fluorescence, an indication that LOV2 reverts to its dark state more quickly than PTP1B. This discrepancy was (i) smallest for PTP1B_PS_, the highest-DR construct, (ii) moderate for 7.1(G528A/N538E), a construct with an intermediary DR, and (iii) largest for 7.1(I532E), a Jα-destabilized mutant without light-dependent catalytic activity (Fig. 2g). This pattern in recovery kinetics provides direct evidence that strong inter-domain conformational coupling is necessary for photocontrol of PTP1B activity.

### Preparation of a Natively Localized Variant of PTP1B_PS_

Inside the cell, PTP1B possesses a C-terminal region—a disordered proline-rich domain followed by a short membrane anchor—that localizes it to the endoplasmic reticulum (ER; Fig. 3a)^34^. To examine the influence of this region on photocontrol, we attached the bulk of it (all but the hydrophobic ER anchor) to the C-terminus of PTP1B_PS_ and assayed the extended chimera *in vitro*. This construct (PTP1B_PS*_) exhibited a reduced DR, which was not improved by the addition of stabilizing mutations to the Jα helix (Figs. 3b-3c); nonetheless, it remained photoswitchable. A construct with the full-length C-terminus of PTP1B (everything including the ER anchor; PTP1B_PS**_), in turn, conferred native localization in COS-7 cells (Fig. 3d).

**Figure 3.**
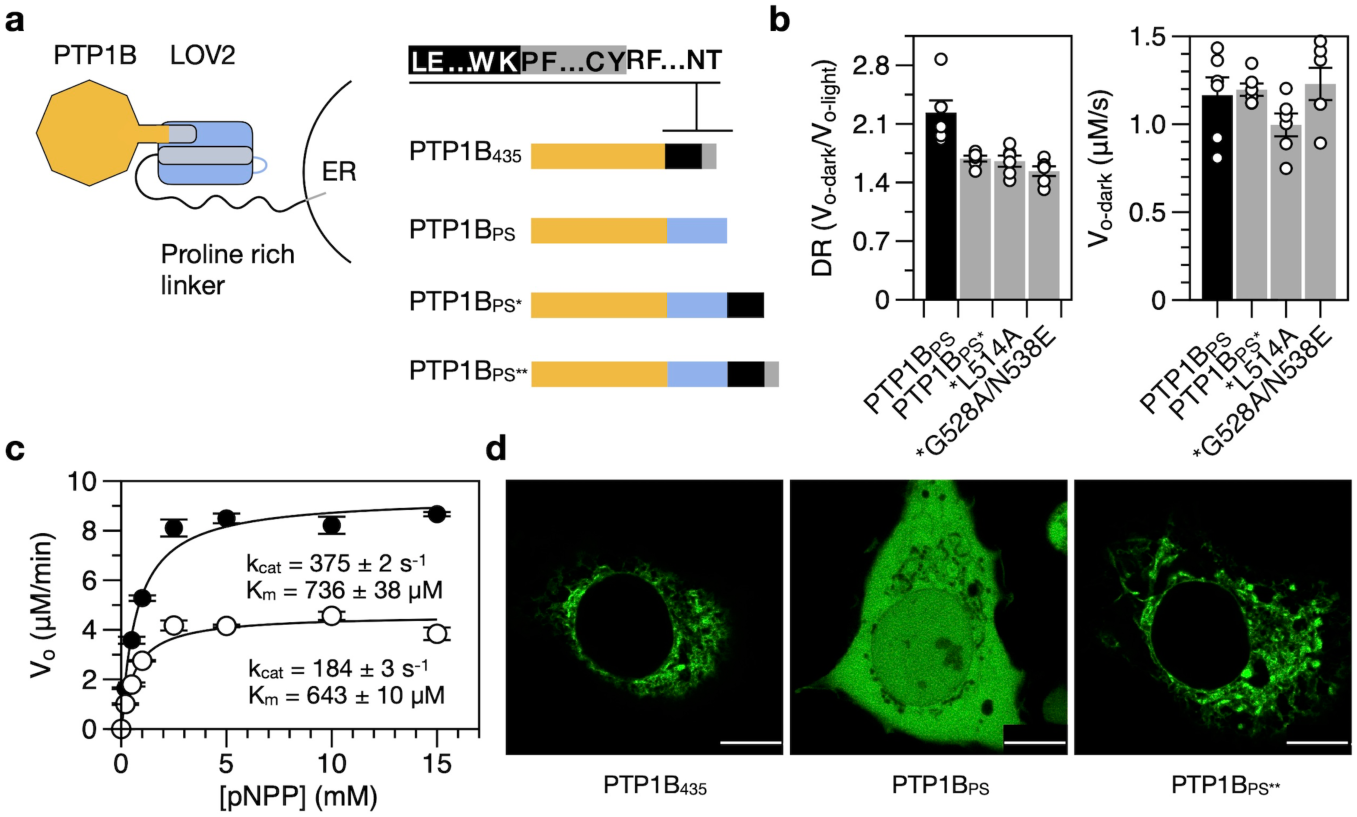
A natively localized variant. **a**, We assembled PTP1B_PS*_ and PTP1B_PS**_ by attaching residues 299-405 (the disordered proline-rich region) and 299-435 (the proline-rich region and ER anchor), respectively, of full-length PTP1B to the C-terminus of PTP1B_PS._ Colors correspond to the catalytic domain of PTP1B (orange), the LOV2 domain (blue), the proline-rich region of PTP1B (black), and the ER anchor of PTP1B (gray). **b**, PTP1B_PS_* is photoswitchable but exhibits a reduced DR, relative to PTP1B_PS_; mutations that stabilize the Jα helix do not improve DR. The plotted data depict the mean, SE, and associated estimates of DR for n = 6 independent experiments. **c,** Saturation curves show the activity of PTP1B_PS*_ on pNPP (*k_cat-dark_*/*k_cat-light_* = 2.03 +/- 0.04). Error bars denote SE for n = 3 independent reactions. **d**, Images of COS-7 cells expressing GFP-tagged variants of PTP1B. PTP1B_435_ and PTP1B_PS**_ exhibit indistinguishable localization patterns (scale bars appear as a white line over a small black rectangle in the lower right corner of each image;10 μm). Source data are provided as a Source Data file.

We completed our characterization of the elongated forms of PTP1B_PS_ by examining the influence of LOV2 on interactions mediated by the disordered C-terminal region. Briefly, we compared the susceptibilities of PTP1B_1-405_ and PTP1B_PS*_ to inhibition by DPM-1001, an inhibitor that binds preferentially to this region (Supplementary Figure 5)^35^. To our surprise, IC_50_’s differed by ∼30%; this small difference indicates that LOV2 does not preclude regulatory interactions with the disordered region. Intriguingly, DPM-1001 also binds weakly to the catalytic domain, likely by binding near the α7 helix^36^; IC_50_’s for PTP1B_1-321_ and PTP1B_PS_ were, thus, much higher than IC_50_’s for the full-length constructs and exhibited a greater sensitivity to LOV2. The light-sensitive domain may thus, affect weak interactions that occur at its point of attachment (though, this region is not an established target of post-translational modifications).

### An Optogenetic Probe for Studying Intracellular Signaling

To examine the function of PTP1B-LOV2 chimeras in living cells, we sought a genetically encodable sensor for PTP1B activity. Several previously developed sensors for PTKs could plausibly support such a function; we chose a sensor for Src kinase^37^, an enzyme with an orthogonal activity to PTP1B^38^. This biosensor consists of an SH2 domain, a flexible linker, and a substrate domain (i.e., WMEDYDYVHLQG, a peptide derived from p130cas), all sandwiched between two fluorescent proteins (FPs). Src-mediated phosphorylation of the substrate domain causes it to bind to the SH2 domain, reducing Förster resonance energy transfer between the FPs (FRET; Fig. 4a); PTP1B-mediated dephosphorylation of the substrate domain, in turn, reverses this effect and increases FRET. To enhance the compatibility of the sensor with the blue light necessary to stimulate LOV2, we replaced CFP and YPet—the original FPs—with mClover3 and mRuby3, which have longer excitation wavelengths^39^. As expected, incubation of the modified sensor with Src reduced FRET and increased the donor/acceptor emission ratio; simultaneous incubation with Src and PTP1B (or Src and EDTA), by contrast, prevented this increase (Figs. 4b).

**Figure 4.**
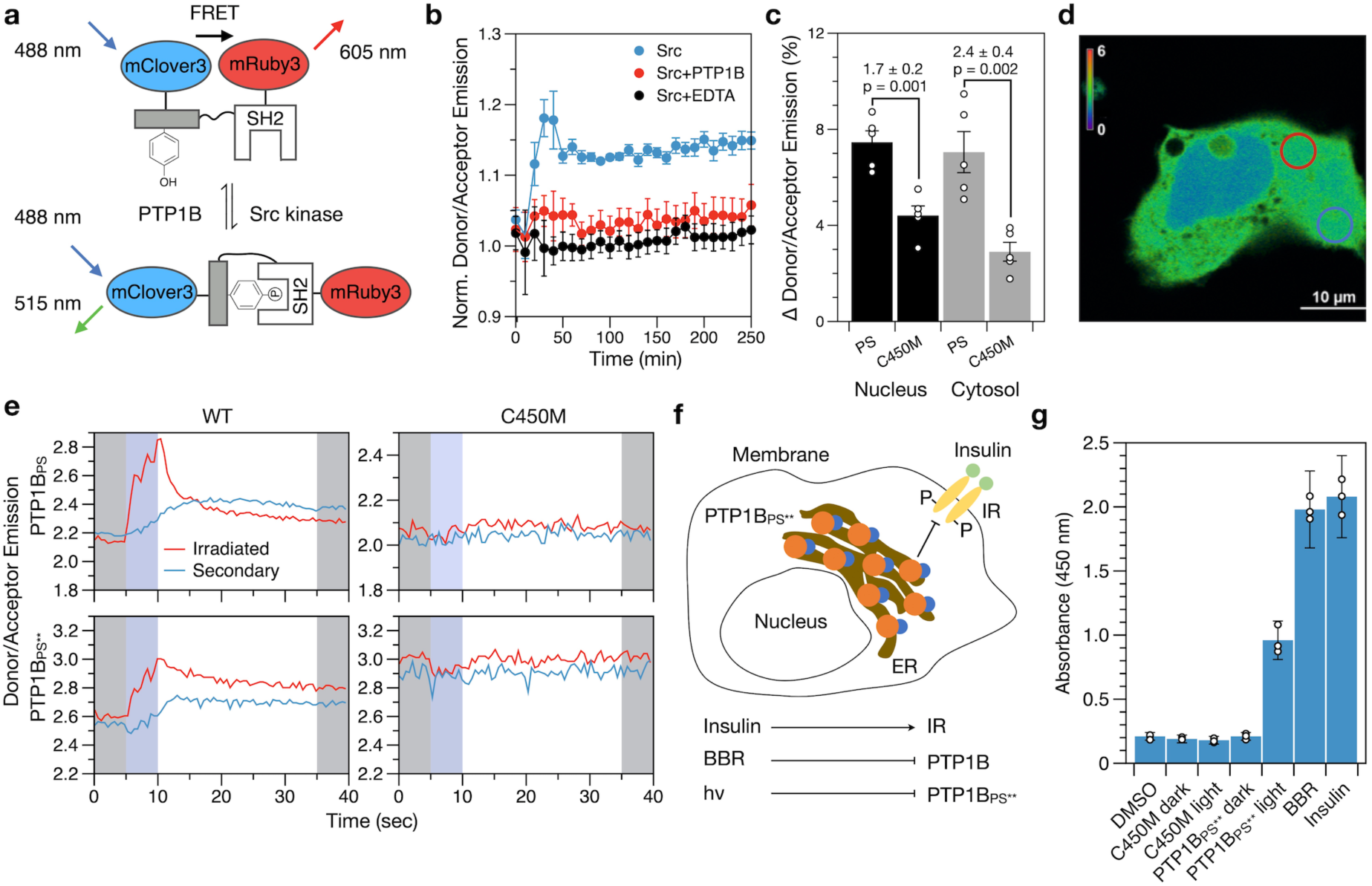
Localized deactivation of PTP1B_PS_. **a,** A biosensor for PTP1B activity. Src-mediated phosphorylation of the substrate domain causes it to bind SH2, triggering a conformational change that decreases FRET; dephosphorylation by PTP1B increases FRET. **b,** Src increases the donor/acceptor emission ratio *in vitro* (normalized by the buffer-only condition); EDTA or PTP1B prevent this increase. Error bars denote propagated SE for measurements of n = 3 independent experiments (measurements are normalized to a to a buffer-only condition). **c,** The percent change in donor/acceptor emission ratio over 1 min within 5-μm circular regions located in the cytosol and nucleus of COS-7 cells activated with 457 nm light. Each condition includes the interquartile average, associated data points, and SE for n = 11 biological replicates. The p-values correspond to a two-tailed Student’s t test. **d**, An image of localized illumination (405 nm) of a COS-7 cell expressing both PTP1B_PS_ and biosensor. Circles delineate irradiated (red) and secondary (blue) regions. **e**, Time courses of FRET in irradiated and secondary regions. Shading highlights 5-s periods before (gray), during (blue), and after (gray) illumination. **f**, A depiction of a HEK293T/17 cell expressing PTP1B_PS**_. Insulin stimulates phosphorylation of the membrane-bound insulin receptor (IR); PTP1B dephosphorylates it. **g**, ELISA-based measurements of IR phosphorylation in (i) wild-type HEK293T/17 cells and (ii) HEK293T/17 cells stably expressing PTP1B_PS**_ or PTP1B_PS**_(C450M). Insulin-mediated simulation of IR, BBR-mediated inhibition of PTP1B, and photoinactivation of PTP1B all increase IR phosphorylation. The dark state of PTP1B_PS**_ and the dark and light states of PTP1B_PS**_(C450M), by contrast, leave IR phosphorylation unaltered from its levels in the wild-type strain (DMSO). The plotted data depict the mean, propagated SE, and associated data points for measurements of n = 3 biological replicates (relative to a buffer-only condition). Source data are provided as a Source Data file.

We began our imaging studies by co-expressing the biosensor with PTP1B-LOV2 chimeras in COS-7 cells. These cells are large and flat and, thus, facilitate imaging of subcellular regions^40^; previous studies have used them to examine PTP1B-mediated signaling events^41, 42^. Whole-cell irradiation of cells expressing PTP1B_PS_ with 457 nm light increased the biosensor signal in both the nucleus and cytosol by ∼7%, a change larger than the 3-4% increase afforded by the dark-state mutant (Fig. 4c and Supplementary Figure 6); the response of the biosensor in cells expressing PTP1B_PS**_, by contrast, was nearly imperceptible when compared to the dark-state analogue (Supplementary Figure 7). We note: Our imaging experiments rely on basal Src activity (i.e., we do not overexpress this enzyme). Accordingly, our findings indicate that transient inactivation of PTP1B allows background concentrations of Src to effect a rapid increase in the population of phosphorylated biosensor (other kinases could certainly contribute to this response, but our chosen biosensor is fairly specific for Src^43^).

Local irradiation of photoswitchable enzymes can permit detailed studies of spatially dependent signaling events and, by minimizing cellular exposure to optical stimuli, reduce the background signal caused by photobleaching^10^. To assess the compatibility of our light-sensitive chimeras with spatiotemporal studies, we used 405-nm light to irradiate 5-µm circular regions within COS-7 cells, and we measured the response of the biosensor (Figs. 4d-4e). In cells expressing PTP1B_PS_, local irradiation of the cytosol produced a transient spike in donor/acceptor emission ratio within the irradiated region and a modest, smooth increase in signal within a secondary region located far from the first (Fig. 4e and Supplementary Figure 8); both irradiated and secondary regions maintained a similar increase in signal for at least 30 seconds after irradiation. In cells expressing PTP1B_PS**_, irradiation near the nucleus produced a similar change in signal, while irradiation near the plasma membrane (PM) failed to do so (Fig. 4e and Supplementary Figures 9-11a). In all cases, dark-state mutants produced no detectable effect. Our results thus indicate that localized inactivation of PTP1B_PS_ and nucleus-proximal PTP1B_PS**_ can produce a measurable cell-wide increase in the phosphorylation state of their targets.

The ER is a vesicular network that extends unevenly from the nucleus of the cell. To determine if the reduced activity of PTP1B_PS**_ near the PM results from the low abundance of ER in this region, we used BFP-Sec61β, a genetically encoded ER label^44^, to measure the subcellular distribution of ER. As expected, the fluorescence of 5-µm circular regions located near the PM was 2.7-fold lower than the fluorescence of equivalently sized regions located near the nucleus (Supplementary Figures 11b-11c); this discrepancy suggests that the diffuse distribution of PTP1B_PS**_ near the PM limits its activity on membrane-proximal targets.

Cells rely on complex networks of biomolecular interactions to transmit, filter, and integrate chemical signals^45^. The biochemical repercussions of changes in the activity of any single regulatory enzyme are, thus, difficult to assess with artificial biosensors alone. To evaluate the influence of modest changes in PTP1B activity on the phosphorylation state of a native regulatory target, we generated HEK293T/17 cells that stably express PTP1B_PS**_ and used an enzyme-linked immunosorbent assay (ELISA) to measure shifts in IR phosphorylation caused by transient illumination (455 nm, 10 minutes). To our surprise, illumination increased IR phosphorylation to levels that rivaled those elicited by high concentrations of both an allosteric inhibitor (BBR) and insulin (Fig. 4g). These optically derived shifts—which, by our best estimate, exceed a twenty-fold increase over background levels (Supplementary Note 2)—did not occur in cells shielded from light or in cells stably expressing PTP1B_PS**_(C450M); IR phosphorylation levels in these two varieties of cells were indistinguishable from those of the wild-type cell line. Our findings thus suggest that PTP1B_PS**_ leaves native phosphorylation levels intact but enables large shifts in target phosphorylation under blue light.

## DISCUSSION

The study of PTPs has long suffered from a paucity of tools for probing and measuring their intracellular activities^46^. In this study, we developed a photoswitchable variant of PTP1B and used it to exert spatiotemporal control over the phosphorylation state of a genetically encoded biosensor in living cells. Transient irradiation of the full-length, natively localized construct near the nucleus but not the PM produced cell-wide changes in sensor phosphorylation. Importantly, the changes in activity afforded by our allosteric control system reach—by our best estimate—70-85% of the maximum achievable dynamic range and match physiologically influential changes in activity caused by post-translational modifications of PTP1B inside the cell (Supplementary Table 1). Our analysis of IR phosphorylation, in turn, suggests that modest changes in the activity of PTP1B are sufficient to effect large shifts (i.e., over twenty-fold) in the phosphorylated fraction of its regulatory targets. This result, which evidences an intriguing sort of hypersensitivity, indicates that our photoswitchable PTP1B—which unlike PTP inhibitors^47^, offers both exquisite selectivity and spatiotemporal precision—could provide a useful tool for biochemical analyses that rely on precise, protein-specific perturbations (e.g., imaging studies and proteomic analyses)^48^. PTP1B is a therapeutic target for the treatment of diabetes, obesity^49^, breast cancer^50^, and cardiovascular disease^51^, and has emerged as a potential modulator of inflammation^52^, anxiety^53^, immunity^54^, memory^55^, and neural specification in embryonic stem cells^56^; by facilitating detailed analyses of the contribution of PTP1B to these complex processes, the tools developed in this study could help to elucidate the biochemical basis—and, perhaps, shared origin—of a diverse set of physiological states.

Classical—or tyrosine-specific—PTPs possess several features that are particularly incompatible with conventional approaches to optical control: Their solvent-exposed active sites are distal to both termini and, thus, difficult to obstruct with light-sensitive fusion partners ^57^; they engage in protein-protein interactions at delocalized—and incompletely mapped—surface sites that make the physiological repercussions of domain insertion (or domain dissection) difficult to assess^58–60^; and their subcellular localization affects regulatory function in a non-binary manner that complicates the use of optically induced re-localization to study cell signaling^41, 61^. Accordingly, future efforts to use these methods to build photoswitchable PTPs could be worthwhile—and may yield constructs with higher DRs than those described in this study; accompanying analyses of the influence of new control systems on protein structure, activity, and/or subcellular localization, however, will facilitate a detailed assessment of their benefits over the present system.

The results of this work offer two general insights for the design of genetically encoded probes. First, kinetic assays of PTP1B_PS_ demonstrate that allosteric systems for photocontrol can—at least, within some classes of proteins—effect isolated changes in *k_cat_*; such designs may be less disruptive to substrate binding (i.e., more substrate agnostic) than systems in which light-sensitive domains act as competitive inhibitors (which can be outcompeted by a sufficiently high concentrations of substrate). Future analyses of differences in the intracellular concentrations and binding affinities of regulatory targets will clarify the extent of this benefit. Second, spectroscopic analyses of PTP1B-LOV2 chimeras provide experimental evidence that strong interdomain conformational coupling can enhance allostery-derived photocontrol of one domain with another. Computational methods to optimize this coupling (e.g., methods that enhance correlated motions between fused domains or that reduce the dissipation of energy between them) could, thus, facilitate the development of new—and, perhaps, protein-specific—varieties of minimally disruptive control systems.

## METHODS

### Cloning and molecular biology

We constructed PTP1B-LOV2 chimeras by fusing PTP1B and LOV2 at crossover points in a primary sequence alignment. In brief, we used EMBOSS Needle, an implementation of the Needleman-Wunsch algorithm^62^, to align the C-terminus of PTP1B (residues 285-305) with the N-terminus of LOV2 (residues 387-410), and we selected eight matching aligned residues as fusion points for the two domains (Fig. 1b). To assemble chimeric genes, we amplified DNA encoding PTP1B and LOV2 from pET21b and pTriEx-PA-Rac1 plasmids, respectively. (The pET21b plasmid was a kind gift from the Tonks Group of Cold Spring Harbor Laboratory; we purchased pTriEx-PA-Rac1 from Addgene). We joined the two amplified segments with overlap extension PCR (oePCR; see Supplementary Table 3 for primers) and ligated the final chimeric product into pET16b for protein expression.

We generated additional constructs with standard techniques. To build single-site mutants and truncation variants, we amplified parent plasmids with appropriate mutagenic primers (Supplementary Table 4). To construct PTP1B_PS*_ and PTP1B_PS**_, we amplified C-terminal regions of PTP1B (residues 299-405 and 299-435, respectively) from pGEX-2T-PTP1B (Addgene) and used Gibson assembly to join them to the C-terminus of PTP1B_PS_ (50°C for 1 hr; see Supplementary Table 5 for primers). Finally, to construct GFP-tagged versions of PTP1B_PS_, PTP1B_PS**_, and PTP1B_435_, we amplified these genes from their parent plasmids (see Supplementary Table 5 for primers) and ligated the PCR product into pAcGFP1-C1 (Clonetech, Inc.) at the NcoI and BamHI sites of the MCS for protein expression.

We developed a biosensor for PTP1B by replacing the fluorescent proteins of a biosensor for Src kinase^37^ with mClover3 and mRuby3^39^. In brief, we amplified DNA encoding the following components: (i) the central segment of the Src biosensor—the SH2 domain, interdomain linker, and substrate domain (i.e., WMEDYDYVHLQG, a peptide derived from p130cas)—from its parent plasmid (a Kras-Src FRET biosensor, Addgene), (ii) genes for mClover3 and mRuby3 (plasmids pNCS-mClover3 and pNCS-mRuby3, respectively), and (iii) the backbone of pAcGFP1-C1 (Clonetech, Inc.). After amplification, we joined all segments with Gibson assembly (50°C for 1 hr; see Supplementary Table 6 for primers).

For live-cell studies, we integrated the modified biosensor and PTP1B-LOV2 chimeras into pAcGFP1-C1 by using protocols described above. In short, we amplified DNA encoding (i) PTP1B_PS_ or PTP1B_PS**_, (ii) a ribosomal skipping peptide sequence (P2A-GSG, GSGATNFSLLKQAGDVEENPGP), (iii) the modified biosensor, and (iv) the pAcGFP1-C1 backbone, and we joined the segments with Gibson assembly (50°C for 1 hr; see Supplementary Tables 6 and 7 for primers and DNA fragments, respectively).

### Protein expression and purification

We overexpressed PTP1B_1-281_, PTP1B_1-321_, PTP1B_1-405_, LOV2_404-547-_, PTP-LOV2 chimeras, Src_251-536_, and the modified biosensor in *E. coli* by carrying out the following steps: (i) We subcloned 6x polyhistidine-tagged versions of each construct into a pET16b plasmid. We positioned the tag at the N-terminus of Src and the FRET-based biosensor and the C-terminus for all other proteins. For Src, we also added a gene for Cdc37, a chaperone that facilitates protein folding in bacteria^63^. (ii) We transformed *E. coli* BL21(DE3) cells (New England Biolabs C2527) with each plasmid and spread the transformed cells onto an agar plate (25 g/L LB, 100 mg/L carbenicillin, 1.5% agar). (iii) We used one colony from each plate to inoculate a 20-mL culture (25 g/L LB and 100 mg/L carbenicillin), which we incubated in a shaker at 37°C overnight. (iv) We used the overnight culture to inoculate 1 L of induction media (20 g/L tryptone, 10 g/L yeast extract, 5 g/L NaCl, 4 g/L M9 slats, 4 g/L glucose, and 100 mg/L carbenicillin), which we incubated in a shaker at 37°C until it reached an OD_600_ of ∼0.6. (v) We induced protein expression by adding 100 μL of 1 M solution of isopropyl β-D-1-thiogalactopyranoside (IPTG) to each culture and by reducing the temperature to 22°C. (vi) At 7 h, we pelleted cells (3950 xg, 20 min; JA-10 Beckman Coulter).

We purified all proteins with fast protein liquid chromatography. To begin, we lysed cell pellets by adding the following components to each gram of pellet: 4 mL of B-PER (Thermo Fisher Scientific, Inc.), 1 mg MgSO_4_, 2 mg Nα-p-Tosyl-L-arginine methyl ester hydrochloride, 1.25 mg tris(2-carboxyethyl)phosphine (TCEP), 3.75 µl phenylmethylsulfonyl fluoride, 1 mg Lysozyme, and 10 µl DNase. After mixing to homogeneity, we rocked the lysis mixtures for 1 h at room temperature (∼22°C), pelleted the cell debris (3950 xg, 60 min), and isolated the supernatant. To clarify the supernatant further, we added a saturated solution of ammonium sulfate to 10% (v/v), pelleted the resulting mixture (3950 xg, 15 min), and used a 0.22-μm filter to remove particulates. To begin purification, we exchanged the filtered supernatant into Tris-HCl buffer (50 mM Tris-HCl, 0.5 mM TCEP, pH 7.5), flowed the exchanged solution over an Ni column, and eluted the protein of interest with a 0-100% gradient of imidazole (50 mM Tris-HCl, 0.5 mM TCEP, 500 mM imidazole, pH 7.5). For further purification, we exchanged each protein into HEPES buffer (50 mM HEPES, 0.5 mM TCEP, pH 7.5), flowed the exchanged solution over an anion exchange column, and eluted the final protein with 0-100% gradient of NaCl (50 mM HEPES, 0.5 mM TCEP, 500 mM NaCl, pH 7.5). We purchased all columns (26/10 HiPrep [desalting], HisTrap HP [Ni], and HiPrep Q HP 16/10 [anion exchange]) from GE Healthcare, Inc. We confirmed the purity of final solutions with SDS-PAGE, and we stored each protein in storage buffer (50 mM HEPES, 0.5 mM TCEP, 20 v/v% glycerol, pH 7.5) at −80°C.

### Initial analysis of photoswitching

We screened PTP-LOV2 chimeras for light-dependent catalytic activity by measuring their activity on 4MUP in the presence and absence of light. In brief, we carried out the following steps: (i) In a room illuminated with a red light (625 nm), we prepared two 96-well plates—hereafter referred to as the “light plate” and “dark plate”—with 100-μL reactions consisting of buffer (50 mM HEPES, 0.5 mM TCEP, pH 7.5), substrate (500 μM 4MUP), and enzyme (5 nM); we added a plastic cover to each plate. (ii) We encased the dark plate in foil and placed the light plate in a chamber made up of two opposing reflective steel bowls fed with a 455-nm light (∼450 mW, SLS-0301-C, Mightex Systems, Inc.; Supplementary Figures 1d-1f). (iii) We incubated both plates at room temperature (∼22°C). (iv) At 7, 14, 21, 28, 35 and 42 minutes after beginning the reaction, we removed each plate from its resting position (i.e., the foil cover or light chamber), loaded it into a SpectraMax M2 plate reader, and measured the formation of 4-methylumbelliferone (365_ex_/450_em_); we immediately returned each plate to its resting position. (v) We used discrete measurements to estimate initial rates and, thus, to calculate DR (i.e., V_o-dark_/V_o-light_).

We minimized error in our measurements of photoswitching with four precautions: (i) We used concentrations of enzyme and substrate that sustained initial reaction rates for 42 minutes, a length of time that minimizes the disruption of 1-min breaks required to measure product formation. (ii) For each construct in each plate, we prepared two sets of three compositionally identical, yet differentially positioned wells; this arrangement minimizes potential contributions from nonuniform illumination. (iii) For each construct at each illumination condition, we repeated the assay at least three times, collecting a total six estimates of initial rate, each based on measurements from three wells. (iv) We established a control range: When wild-type PTP1B, which was present in each plate, exhibited a 10% difference in activity between the two plates, we discarded data from both (i.e., we assumed that differences in activity between the two plates were not caused by the presence or absence of light).

We examined the light-dependent catalytic activity of PTP1B_PS_ on a phosphopeptide (DADEpYLIPQQG from EGFR) by following the aforementioned procedure with several differences: (i) We used a substrate concentration of 120 µM and a total reaction volume of 40 uL. (ii) We added malachite green solution (Sigma-Aldrich) to stop individual reactions at 2, 4, 6, or 8 minutes. (iii) We measured the formation of phosphate by using the plate reader to quantify a complex formed between orthophosphate, molybdate, and Malachite Green (620_abs_); we waited until the end of our experiments (i.e., 8 minutes) for all absorbance measurements. We note: the statistically indistinguishable DRs (p < 0.01, two-tailed Student’s t test) determined for PTP1B_PS_ on substrates that require different spectrophotometric measurements (i.e., fluorescence at 450 nm for 4MUP and absorbance at 620 nm for a phosphopeptide) suggest that the illumination conditions used for optical measurement do not artificially depress or enhance DR.

### Enzyme kinetics

We examined the influence of photomodulation on enzyme kinetics by measuring the activities of PTP1B_PS_ and PTP1B_PS*_ on pNPP in the presence and absence of light (i.e., we used dark and light plates as described above). Briefly, we prepared 100-μL reactions consisting of buffer (50 mM HEPES, 0.5 mM TCEP, pH 7.5), substrate (0.2, 0.5, 1, 2.5, 5, 10 and 15 mM pNPP), and enzyme (25 nM); at 4, 8, 12, 16 and 20 minutes after initiating the reaction, we measured the production of p-nitrophenol (405_abs_) on a SpectraMax M2 plate reader; and we used DataGraph to fit initial rates to a Michaelis-Menten model of enzyme kinetics. Final values of *k_cat_* and *K_m_* reflect the mean of independent estimates determined from three Michaelis-Menten curves; error *k_cat_* and *K_m_* reflects the standard error of those estimates.

We examined the inhibitory effect of DPM-1001 on PTP1B-mediated hydrolysis of pNPP as follows: (i) We carried out the aforementioned pNPP reactions in the presence of different concentrations of DPM-1001 (0, 20, 40, 60 μM for PTP1B_1-405_ and PTP1B_PS*_; 0, 100, 200, 400 μM for PTP1B_321_ and PTP1B_PS_). (ii) We used MATLAB’s “nlinfit” and “fminsearch” functions to fit (a) initial-rate measurements collected in the absence of inhibitors to a Michaelis−Menten model and (b) initial-rate measurements collected in the presence and absence of inhibitors to four models of inhibition (i.e., competitive, noncompetitive, uncompetitive, and mixed inhibition^64^). (iii) We used an F-test to compare the fits of (a) a mixed model, which has two parameters, and (b) each nested single-parameter model with the lowest sum of squared errors for a given dataset. DPM-1001 exhibited mixed inhibition for all constructs (p < 0.01, two-tailed Student’s t test). (iv) We estimated IC_50_’s by using the best-fit kinetic model to determine the inhibitor concentration required to reduce initial rates by 50% on 15 mM pNPP. This high substrate concentration minimizes the concentration dependence of IC_50_’s. We used the MATLAB function “nlparci” to determine the confidence intervals of kinetic parameters and propagated those intervals to estimate the corresponding confidence on IC_50_’s.

We compared the activities of PTP1B_1-281_ and PTP1B_1-321_ on pNPP by using a continuous assay. Briefly, we prepared 100-μL reactions consisting of buffer (50 mM HEPES, 0.5 mM TCEP, pH 7.5), pNPP (0.2, 0.5, 1, 2.5, 5, 10 and 15 mM), and enzyme (25 nM); we measured the production of p-nitrophenol at five-second intervals for 270 seconds (SpectraMax M2 plate reader); and we used DataGraph to fit initial rates to a Michaelis-Menten model. Final values of *k_cat_* and *K_m_* reflect the mean of independent estimates determined from three Michaelis-Menten curves; error *k_cat_* and *K_m_* reflects the standard error of those estimates.

Finally, we evaluated the reversibility of our LOV2-based light switch by illuminating 25 uM of PTP1B_PS_ (50 mM HEPES, 0.5 mM TCEP, pH 7.5) for 10 seconds and by, subsequently, monitoring its activity on 5 mM pNPP after 5 minutes in the dark. To minimize error, we repeated this experiment three times with seven cycles per experiment (Supplementary Figure 2).

### X-ray crystallography

We prepared crystals of PTP1B_PS_ by using hanging drop vapor diffusion. To begin, we prepared a concentrated solution of PTP1B_PS_ (∼400 μM PTP1B_PS_, 50 mM HEPES, pH 7.3) and a crystallization solution (100 mM HEPES, 200 mM magnesium acetate, and 14% polyethylene glycol 8000, pH 7.5); we mixed the two solutions in 1:2, 1:3, and 1:6 ratios (protein: crystallization) to form 7-9 μl droplets for crystal growth; and we incubated the droplets over reservoirs filled with crystallization solution at 4℃ in the dark. Long hexagonal crystals with a yellow hue appeared after 1-3 weeks. Prior to freezing, we soaked all crystals in cryoprotectant (100 mM HEPES, 200 mM magnesium acetate, and 25% polyethylene glycol 8000, pH 7.5) overnight.

We collected X-ray diffraction through the Collaborative Crystallography Program of the Berkeley Center for Structural Biology. We performed integration, scaling, and merging of XRD data with the xia2 software package, and we carried out molecular replacement with the Phenix graphical user interface, followed by one round of PDB-REDO^65^. The crystallographic data collected in this study are reported in Table 1.

**Table 1.** Data collection and refinement statistics for X-ray crystallographic analysis of PTP1B_PS_

|  | PTP1B<sub>PS</sub> (6NTP) |
| --- | --- |
| **Data collection** |  |
| Space group | P3<sub>1</sub>21 |
| Cell dimensions |  |
| *a*, *b*, *c* (Å) | 89.254, 89.254, 105.747 |
| $\alpha$, $\beta$, $\gamma$ (°) | 90.000, 90.000, 120.000 |
| Resolution (Å)<sup>a</sup> | 52.87 – 1.89 (1.92-1.89) |
| *R*<sub>merge</sub><sup>a</sup> | 0.071 (2.550) |
| *R*<sub>pim</sub><sup>a</sup> | 0.025 (0.893) |
| <*I*/$\sigma$(*I*)><sup>a</sup> | 17.2 (1.1) |
| CC<sub>1/2</sub><sup>a</sup> | 1.000 (0.626) |
| Completeness (%)<sup>a</sup> | 100.0 (100.0) |
| Multiplicity<sup>a</sup> | 9.0 (9.1) |
| **Refinement** |  |
| Resolution (Å) | 44.67-1.89 |
| No. reflections | 37270 |
| *R*<sub>work</sub> / *R*<sub>free</sub> | 0.17922/0.21175 |
| No. atoms |  |
| Protein | 2351 |
| Ligand/ion | 1 |
| Water | 224 |
| *B*-factors |  |
| Protein<sup>b</sup> | 21.7 |
| Ligand/ion<sup>b</sup> | 50.8 |
| Water<sup>b</sup> | 47.4 |
| R.m.s. deviations |  |
| Bond lengths (Å) | 0.008 |
| Bond angles (°) | 1.488 |

<sup>a</sup>Values in parentheses are for highest-resolution shell. We examined one crystal.
<sup>b</sup>Values correspond to means of *B*-factors for the indicated atoms.

### Circular dichroism spectroscopy

We examined the influence of photomodulation on the secondary structure of PTP1B-LOV2 chimeras by using a circular dichroism spectrophotometer (Applied Photophysics Chirascan Plus) to measure optically induced changes in α-helical content. To collect full-spectrum measurements, we incubated 0.2 g/L solutions of each chimera (10 mM NaPi, 0.5 mM TCEP, pH 7.5) in a crystal cuvette (0.05-cm path length) for 10 seconds with/without blue light (455 nm) and immediately measured mean residue ellipticity (MRE) at 1-nm increments from 185 to 260 nm. To measure thermal recovery, we began as before, but we measured MRE at 222 nm every 2.5 seconds for 250 seconds in the dark. We normalized the CD data thus gathered with Eq. 1, where *CD_t_*, *CD_0_*, *CD_250_* represent MRE at *t*, 0, and 250

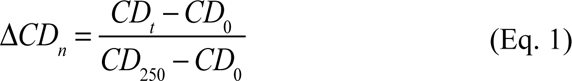

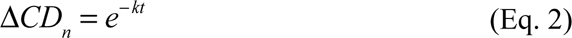

seconds; and we fit the normalized data to an equation for exponential decay (Eq. 2). Final values of *k* reflect the mean and standard error of values determined from fits to six data sets.

### Fluorescence spectroscopy

To examine the influence of photomodulation on the conformation of PTP1B within PTP1B-LOV2 chimeras, we use fluorescence spectroscopy to measure optically induced changes in tryptophan fluorescence. In brief, we prepared 60 µM solutions of protein (50 mM HEPES, 0.5 mM TCEP, pH 7.5) in a Helma ultra-micro quartz cuvette (Thomas Scientific, Inc.); we illuminated those solutions for 10 seconds with a 455-nm light; and we monitored fluorescence (280_ex_/365_em_) in 10-second intervals for 200 seconds using a SpectraMax M2 plate reader. We normalized the fluorescence data, thus gathered, with Eq. 3, where *W_t_, W_0_*,

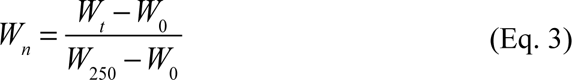

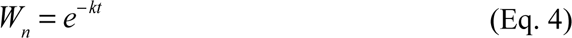

W_250_ represent the emission at *t*, 0, and 250 seconds; and we fit the normalized data to an equation for exponential decay (Eq. 4). Final values of *k* reflect the mean and standard error of values determined from fits to six datasets.

### Biosensor development

We assessed the sensitivity of an Src biosensor^37^ to the activity of PTP1B by incubating it with Src in the presence and absence of PTP1B. We prepared 100-μL reactions consisting of 2 µM biosensor and 300 nM Src kinase in 1X kinase buffer A (Thermo Fisher Scientific, Inc.) supplemented with 2 mM DTT, 2 mM MgCl_2_, and 50 µM ATP. For a subset of reactions, we added PTP1B and EDTA at concentrations of 100 nM and 50 mM, respectively. For each reaction, we monitored the fluorescence of mClover3 (475_ex_/520_em_) and mRuby3 (475_ex_/600_em_ nm) in 10-min increments for 300 minutes on a Spectramax M2 plate reader.

### Preparation of cells for imaging experiments

For live-cell imaging experiments, we grew COS-7 cells (ATCC CRL-1651, seeded from a freezer stock) in DMEM media supplemented with 10% FBS, 100 units/ml penicillin, and 100 units/ml streptomycin for ∼24 hr to achieve 70-90% confluency, and we seeded them on a 20-mm glass-bottom cell culture dish (MatTek). At 10-20 hours after seeding, we depleted endogenous PTP1B by transfecting the cells with 25 nM of a PTP1B siRNA silencer (AM16794, Thermo Fisher Scientific, Inc.), 12.5 μl Dharmafect, and 10% FBS. At 5 hours after adding siRNA, we washed cells with 1X PBS buffer, replaced this buffer with DMEM media supplemented as above (with FBS and antibiotic), and transfected the cells with 2000 ng of plasmid DNA and 6 μl of Lipofectamine 2000 reagent (Invitrogen) according to the manufacturer’s protocols. At 10-12 hour after transfection with plasmid DNA, we imaged the cells in Opti-MEM media at 37°C.

### Confocal microscopy

We carried out all imaging experiments with a 100x 1.45 NA oil objective on a Nikon A1R confocal scanning microscope supplemented with an environmental chamber (37°C, 75% humidity, and 5% CO_2_; Pathology Devices, Inc.). To localize both GFP-tagged PTP1B-LOV2 chimeras and BFP-Sec61β, we illuminated Cos-7 cells with a 488-nm laser (0.57 mW/μm^2^ with a pixel dwell time of 2.2 µs) and imaged them with a 525/50 nm bandpass filter. The plasmid bearing BFP-Sec61β (pTagBFP-C1) was a kind gift from the lab of Gia Voeltz of the University of Colorado, Boulder.

For whole-cell activation studies, we illuminated individual cells with a 457-nm laser focused over the breadth of the cell (0.14 mW/μm^2^ with a pixel dwell time of 4.8 µs). To examine the photoresponse of the biosensor after activation, we illuminated the field of view with a 488 nm laser (0.57 mW/μm^2^) and imaged the entire cell with 525/50 nm and 600/50 nm bandpass filters for 1 minute (resonant scanning mode with 518.1-ms frame time). We estimated the average change in donor/acceptor emission ratio between 0 and 60 seconds after activation by calculating the interquartile average of measurements from 11 individual cells.

For localized activation studies, we focused 405-nm light over 5-µm circular regions (0.49 mW/μm^2^ with a pixel dwell time of 4.8 µs) and imaged the photoresponse of the biosensor by illuminating at 488 nm (0.57 mW/μm^2^; 480/30 nm excitation filter) and imaging with 525/50 nm and 600/50 nm bandpass filters for 1 minute. We estimated the average change in donor/acceptor emission ratio within circular regions, in turn, by calculating the difference in 5-second averages starting (i) 5 seconds before activation and (ii) 35 seconds after activation; final estimates of changes in donor/acceptor emission reflect the mean and standard error from six independent measurements (i.e., six individual cells).

The 488-nm light used to image our FRET-based biosensor could plausibly stimulate LOV2, which absorbs at 488 nm (although less so than at 405 and 457 nm)^66^. The results of Figure 4e, however, indicate that such activation does not occur. In brief, irradiation with 405-nm light causes a transient increase in FRET signal for cells expressing PTP1B_PS_ and PTP1B_PS**_, but not for cells expressing light-insensitive analogues of these two constructs; accordingly, 488-nm light does not activate LOV2 (at least, no fully) under our imaging conditions (if it did so, irradiation at 405 nm would not elicit further activation). The insensitivity of LOV2 to 488-nm light likely results from both (i) the low extinction coefficient of LOV2 at 488 nm and (ii) the insufficient combination of power and pixel dwell time of the 488-nm laser.

### Preparation of cells for enzyme-linked immunosorbent assay (ELISA)

We prepared HEK293T/17 cells (ATCC CRL-11268) stably expressing PTP1B_PS**_ or PTP1B_PS_(C450M) by following standard protocols. In brief, we grew the cells in 75 cm^2^ culture flasks (Corning) with DMEM media supplemented with 10% FBS, 100 units/ml penicillin, and 100 units/ml streptomycin. When cells achieved 60-80% confluency, we transfected them with (i) 2000 ng of plasmid DNA (pAcGFP1-C1 with PTP1B_PS**_ or PTP1B_PS**_(C450M), but no GFP) linearized with the ApaLI restriction enzyme (New England Biolabs) and (ii) 6 μl of Lipofectamine 2000 reagent (Invitrogen) according to the manufacturer’s protocols. We passaged the cells in our growth media (as above) supplemented with 1.5 µg/mL puromycin, and we replaced the media every day for 10 days. We passaged the cells 7 times before freezing them for further use.

### Enzyme-linked immunosorbent assay (ELISA) of insulin receptor phosphorylation

We examined IR phosphorylation in HEK293T/17 cells exposed to various conditions by using an enzyme-linked immunosorbent assay (ELISA). To begin, we used siRNA to deplete cells of endogenous PTP1B (see above) and starved them for 48 hours with FBS-free media. After starvation, we exposed cells to one of several conditions for 10 minutes: (i) 455-nm light (we irradiated the culture flask with ∼450 mW light, SLS-0301-C, Mightex Systems, Inc.), (ii) sustained darkness (i.e., we wrapped the culture flask in aluminum foil), (iii) 300 µM of BBR (3-(3,5-dibromo-4-hydroxybenzoyl)-2-ethyl-N-[4-[(2-thiazolylamino)sulfonyl]phenyl]-6-benzofuransulfonamide, Cayman Chemical), an allosteric inhibitor of PTP1B, (iv) 10 nM human insulin (Sigma), and (v) 1.5% DMSO. After these perturbations, we incubated each sample with lysis buffer (Cell Signaling Technology) supplemented with 1X halt phosphatase inhibitor cocktail and 1X halt protease inhibitor cocktail (Thermo Fisher Scientific) for 10 minutes, spun the cells down, and measured IR phosphorylation by using the PathScan® Phospho-Insulin Receptor β (panTyr) Sandwich ELISA Kit (Cell Signaling Technology; #7082).

We carried out the ELISA by using the manufacturer’s prescribed steps: (i) We diluted the entirety of each lyophilized antibody—a detection antibody (phospho-tyrosine mouse detection mAb; #12982) and a secondary detection antibody (anti-mouse IgG, HRP-linked antibody; #13304)—into 11 ml of antibody-specific diluent (detection antibody diluent; #13339; HRP diluent; #13515). (ii) We used lysis buffer to dilute each sample of cell lysate to 30 mg/ml total protein (based on absorbance at 280 nm). (iii) We prepared 100 µL of 1X, 2X, 4X, and 8X dilutions of lysate from each sample (1X corresponds to no dilution, 2X corresponds to a 1:1 dilution in lysis buffer and cell lysate, and so on), and incubated each 100-µL sample in a single well of an antibody-coated 96-well plate (insulin receptor β rabbit mAb coated microwells; #18872) at 4°C overnight. (iv) We washed the cells four times with 200 µL of 1X wash buffer and incubated the washed cells with 100 µL of detection antibody at 37°C for 1 hr. (v) We washed the cells four times as before and incubated the cells with 100 µL of HRP-linked secondary antibody solution at 37°C for 30 min. (vi) We washed the cells four times and incubated them with 100 µL of TMB substrate at 37°C for 10 min. (vii) We added 100 µL of STOP solution and measured absorbance at 450 nm using SpectraMax M2 plate reader.

### Statistical analysis

We used an F-test to compare one-and two-parameter models of inhibition to one another. For all other analyses, we determined statistical significance by using a two-tailed Student’s t test.

### Reporting summary

Additional information on experimental design is available in the Nature Research Reporting Summary.

### Data availability

The source data underlying Figures 1c, 1e, 1f, 2a, 2b, 2c, 2d, 2e, 2g, 3b, 3c, 4b, 4c, 4e, and 4g and Supplementary Figures 1b, 1c, 4a, 4c, 5b, 5c, 5d, 6b, 7b, 8c, 8d, 9c, 9d, 10c, 10d, 11a, 11c, 12a, and 12b are provided as a Source Data file; this data includes exact sample sizes for each dataset. For clarity, Supplementary Data 1-8 also group data by type (photoswitching experiments, kinetic analyses, FRET-based studies, etc.). The crystal structure determined in this study is available from the RCSB Protein Data Bank (PDB entry 6ntp [https://www.rcsb.org/structure/6ntp]). Table 1 provides the refinement statistics for this structure. Plasmids harboring important genes used in this study are available from Addgene: LOV2 (pTriEx-PA-Rac1, #22024,) full-length PTP1B (pGEX-2T-PTP1B, #8602), and biosensor (Kras-Src FRET biosensor, #78302). All other raw data not included in the manuscript are available from the corresponding author upon request.

## ACKNOWLEDGEMENTS

This work was supported by funds provided by the National Science Foundation (A.H. and J.M.F, award 1804897). The ALS-ENABLE beamlines are supported by the National Institutes of Health (award P30 GM124169-01); the Nikon A1R microscope, by the NIST-CU Cooperative Agreement (award 70NANB15H226). The Advanced Light Source is a user facility sponsored by the Department of Energy’s Office of Science (contract DE-AC02-05CH11231); the Advanced Light Microscopy Core is part of the BioFrontiers Institute. We thank Joe Dragavon for guidance on imaging studies and Annette Erbse for assistance with CD spectroscopy.

## AUTHOR CONTRIBUTIONS

J.M.F. conceived of research. A.H. and J.M.F designed experiments. A.H. carried out cloning, protein expression, kinetic measurements, crystal growth, spectroscopic analyses, and imaging experiments. B.S. collected X-ray diffraction data. B.S. and P.Z. provided guidance on structural refinement. A.H. and J.M.F analyzed all data. A.H. and J.M.F. wrote the paper.

## ADDITIONAL INFORMATION

**Supplementary information** is available in the online version of this paper.

