## Supporting Information for "Minimally disruptive optical control of protein tyrosine phosphatase 1B"

Supplementary Notes 1-2……………..……….……………………………………………….S2

Supplementary Figures 1-12………………………………………………………………… S4

Supplementary Tables 1-10………………………………………………………………….S17

SI References…………………………………………………………………………………...S31

**Supplementary Note 1. Analysis of the crystal structure of PTP1B$_{PS}$.** We used X-ray crystallography to examine the structural integrity of PTP1B within the PTP1B$_{PS}$ chimera (Table 1). Crystals of PTP1B$_{PS}$ exhibited three surprising features (Fig. 1d and Supplementary Figs. 2 and 3): (i) Their unit cell and space group were indistinguishable from those of previously collected crystals of PTP1B and PTP1B-ligand complexes. (ii) They permitted resolution of PTP1B, but not LOV2. (iii) They showed PTP1B with a wild-type conformation (i.e., the root-mean-square deviation of aligned atoms between the catalytic domains of PTP1B$_{PS}$ and wild-type PTP1B was 0.30 Å). These features, considered alone, might indicate that LOV2 is absent from our crystals; four additional crystallographic attributes, however, contradict this interpretation: (i) Our crystals were yellow, a color derived from LOV2 (a consequence of its FMN cofactor[1,2]; Supplementary Figures 2a and 2b). (ii) When exposed to 455-nm light, the crystals turned clear, an indication that LOV2 remains capable of forming a cysteine adduct with FMN (Supplementary Figures 2d and 2e). We note: Solutions of PTP1B$_{PS}$(C450M), which cannot form the cysteine adduct, do not photoswitch (Supplementary Figure 2f). (iii) The unit cell had a gap near the α7 helix, the attachment point of LOV2 (Supplementary Figure 2c). (iv) Previously examined apo structures of PTP1B in which the α7 helix is disordered possess the same unit cell and space group as our crystals.[3] These features, taken together, suggest that disorder in the α7 helix of PTP1B causes variability in the orientation of LOV2 within the crystal lattice. Broadly, PTP1B crystallizes with the same space group, unit cell, and conformation in the presence and absence of LOV2 and, thus, appears to be structurally unperturbed (to the extent detectable with X-ray crystallography) by this light-sensitive fusion partner.

**Supplementary Note 2. Quantitative analysis of insulin receptor phosphorylation with an enzyme-linked immunosorbent assay (ELISA).** The signals afforded by enzyme-linked immunosorbent assays (ELISAs) are notoriously nonlinear[4,5]. To quantify sample-to-sample differences in concentrations of phosphorylated insulin receptor—an analyte the lacks a commercially available standard—we carried out the following steps: (i) We fit a dilution curve (i.e., a plot of assay signal vs. dilution factor) for each sample to a four-parameter logistic equation (Supplementary Table 9). (ii) We used the dilution curve of each sample of interest to estimate the difference in analyte concentration between that "reference" sample and a second sample. For example, on a particular dilution curve (e.g., the curve for HEK293T/17 cells exposed to 10 nM of insulin), the signal afforded by a dilution factor of 1 corresponds to the undiluted analyte concentration in the cell lysate, and the signal afforded by a dilution factor of 0.5 corresponds to one-half of that concentration. By solving for the dilution factor associated with the signal of a second undiluted sample (e.g., PTP1B$_{PS**}$-expressing HEK293T/17 cells exposed to 455 nm light), we estimated the fold-difference in analyte concentration between the two samples (e.g., a dilution factor of 0.25 indicates that the second sample has four-fold less phosphorylated insulin receptor than the first sample). (iii) To check the consistency of our analysis, we used the dilution curve for the second sample to estimate, once again, the difference in analyte concentration between the two samples. In all cases, the two estimates—each based on a separate dilution curve—differed by less than 40%. (iv) We calculated the average and standard deviation the two estimates (Supplementary Table 10 and Supplementary Figure 12). The results of our analysis suggest that transient illumination of PTP1B$_{PS**}$ enables a >20-fold change in the concentration of phosphorylated insulin receptor within the cell.

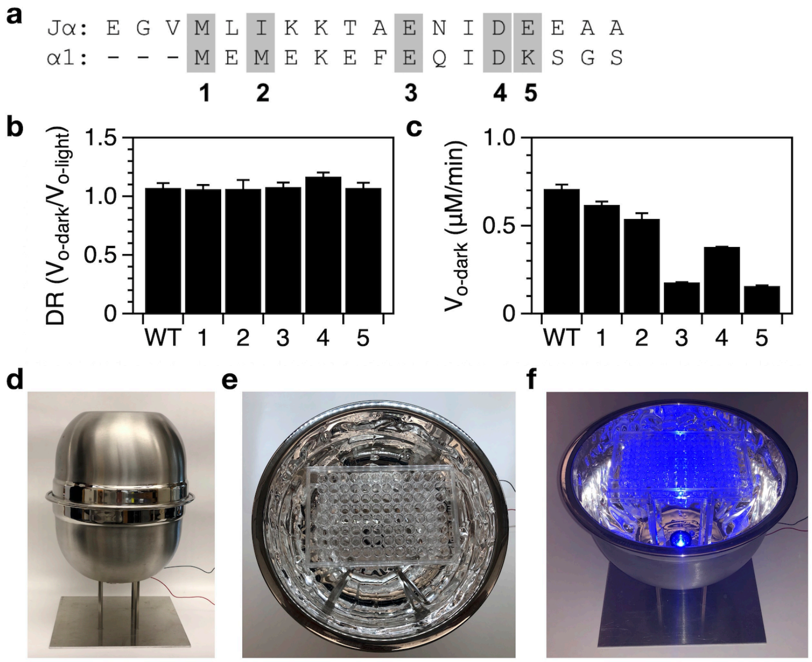


**Supplementary Figure 1 | Analysis of LOV2-PTP1B chimeras. a**, We complemented our initial set of chimeras (Fig. 1b) by constructing a second set in which the C-terminal Jα helix of LOV2 is fused to the N-terminus of PTP1B at homologous crossover points. **b**-**c**, Assays of these chimeras on 4MUP indicate that their catalytic activities are (**b**) light-insensitive and (**c**) significantly reduced (relative to wild-type). The plotted data depict the mean and SE for $n \geq 6$ independent experiments. **d**, The light chamber used for *in vitro* reactions. We coated two stainless steel mixing bowls (Crate and Barrel) with reflective foil and assembled them into a reflective chamber. To the bottom half of this chamber, we added both (i) a 455 nm LED light (SLS-0301-C, Mightex Systems), fed through a hole, and (ii) four steel rods. **e**, The bottom half of the chamber loaded with a 96-well plate. **f**, An image showing the blue light at full power (~450 mW). Source data are provided as a Source Data file.

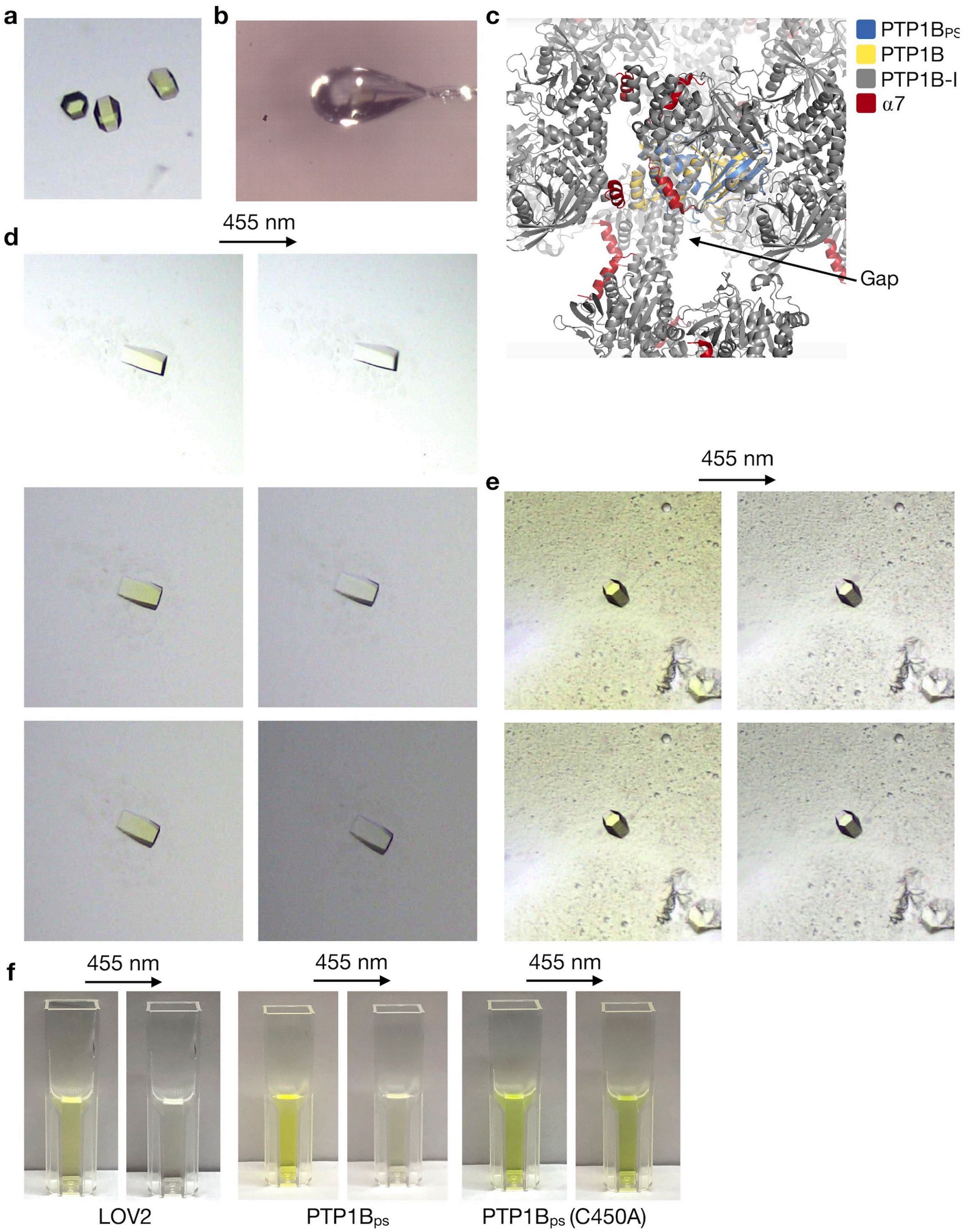


**Supplementary Figure 2 | Crystals of PTP1B-LOV2 chimeras. a**, Crystals of PTP1B-LOV2 chimeras appeared yellow (or yellow green), an indication that they contained LOV2. **b,** The crystal of PTP1B$_{PS}$ used to collect X-ray diffraction data; close inspection reveals a yellow tinge.

**c**, Aligned catalytic domains of PTP1B in three structures: photoswitchable (6ntp), apo (3a5j), and competitively inhibited (2f71). Neighboring asymmetric units from the crystal of the competitively inhibited structure (α7 helix in red) surround the aligned structures (30-Å cutoff). A gap exists near the α7 helix, the attachment point for LOV2. We note: 6ntp and 2f71 have the same space group ($P3_121$) and unit cell dimensions. **d-e,** When exposed to 455-nm light (10-seconds, ~450 mW), crystals of $PTP1B_{PS}$ switched from yellow (oxidized FMN) to clear (reduced FMN); incubation in the dark (5 minutes) restored their yellow color and enabled repeated photoswitching (1x, 2x, and 3x). **f**, When exposed to 455-nm light (10 seconds), solutions containing only LOV2 or $PTP1B_{PS}$ transitioned from yellow to clear, while a solution of $PTP1B_{PS}$(C450M) remained yellow.

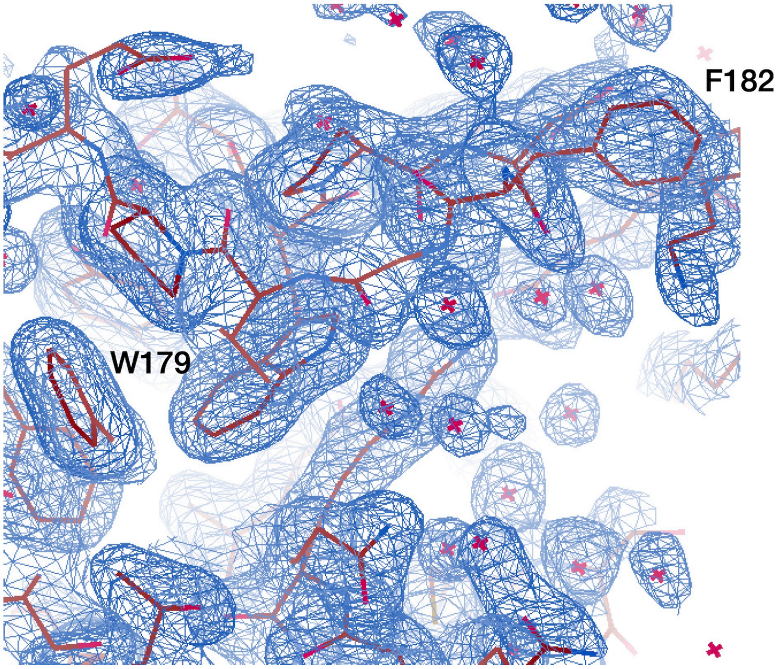


**Supplementary Figure 3 | Crystallographic analysis of PTP1B$_{PS}$.** An electron density map shows the WPD loop of PTP1B in the crystal structure of PTP1B$_{PS}$ (FWT 1.02 σ); the loop is in the open position. As discussed above, the LOV2 domain of PTP1B$_{PS}$ was unresolvable.

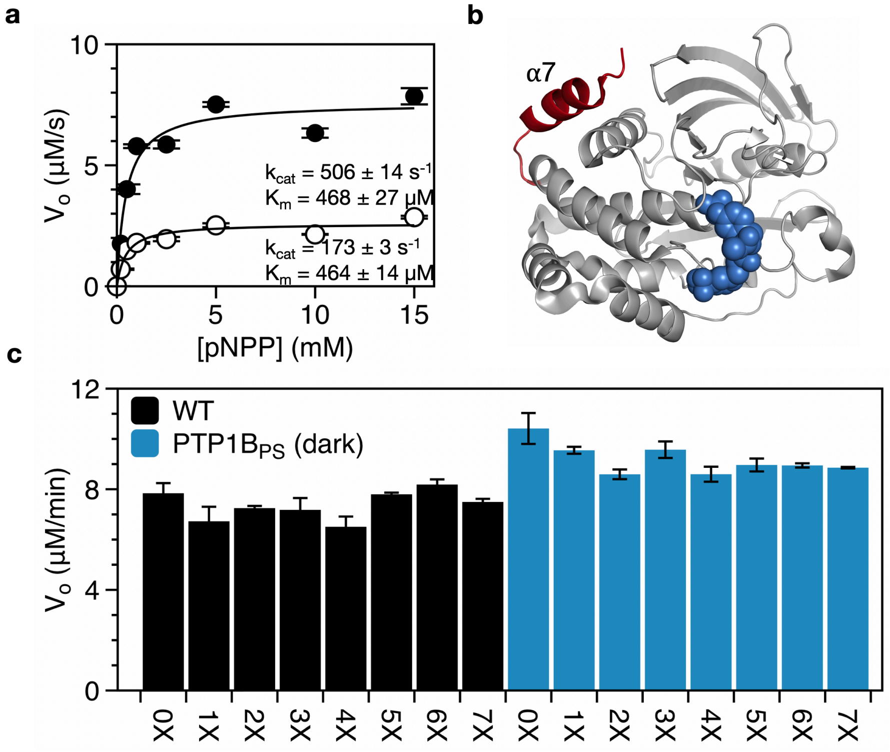


**Supplementary Figure 4 | Analysis of activity on p-nitrophenyl phosphate (pNPP). a**, Initial rates of pNPP hydrolysis by $PTP1B_{321}$ (black) and $PTP1B_{281}$ (white); lines that denote nonlinear least-squares fits to a Michaelis-Menten model. A difference in $k_{cat}$'s of 2.92 ± 0.10 suggests that the DR of $PTP1B_{PS}$ (2.50 ± 0.04 on the same substrate) is 85% of the maximum achievable value for a photoswitch that exploits changes in the conformation of the α7 helix. **b**, A crystal structure of PTP1B (pdb entry 2f71) bound to a competitive inhibitor (blue spheres). Residues 282-298 (red) are missing in $PTP1B_{281}$. **c**, Initial rates of pNPP hydrolysis by $PTP1B_{PS}$ after repeated exposure with 455 nm light (i.e., for each test, we irradiated $PTP1B_{PS}$ with blue light for 10 seconds and waited 5 min after irradiation to perform kinetic assays). $PTP1B_{PS}$ maintains its dark-state activity after multiple cycles of exposure. For **a** and **c,** error bars denote SE for $n \geq 3$ independent reactions. Source data are provided as a Source Data file.

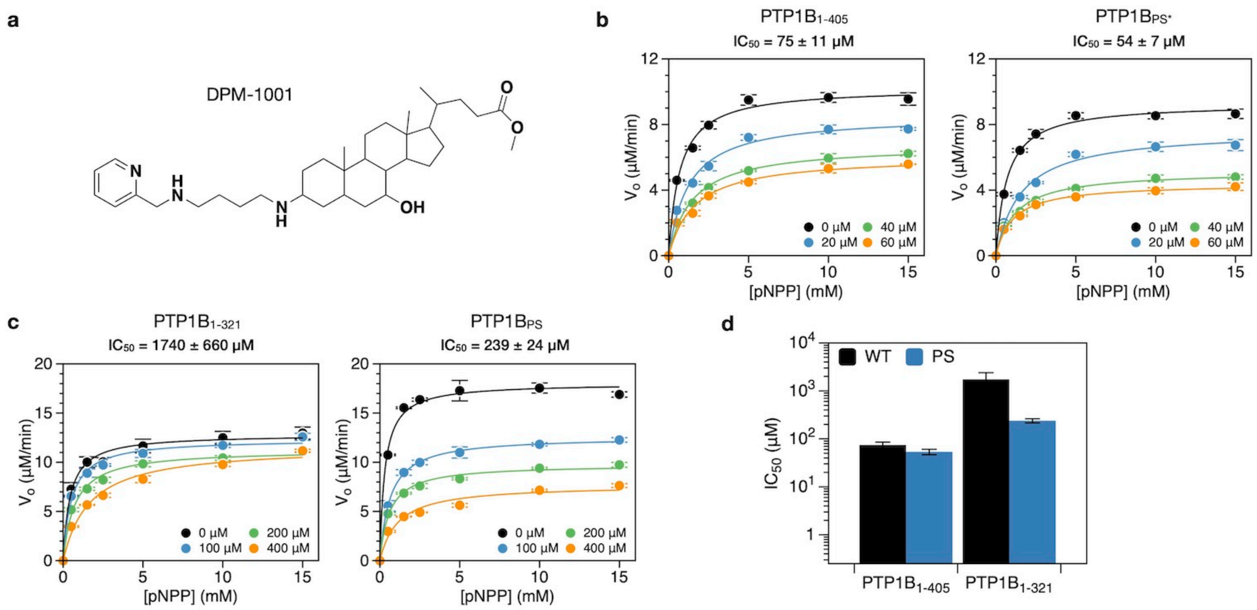


**Supplementary Figure 5 | Analysis of full-length constructs. a**, The molecular structure of DPM-1001, an inhibitor that binds preferentially to the disordered C-terminus of PTP1B. **b**, Inhibition of PTP1B$_{1-405}$ and PTP1B$_{PS*}$ by DPM-1001. **c**, Inhibition of PTP1B$_{1-321}$ and PTP1B$_{PS*}$ by DPM-1001. **d**, A comparison of IC$_{50}$'s for DPM-1001 on wild-type variants (PTP1B$_{1-321}$ and PTP1B$_{1-405}$) and photoswitchable constructs (PTP1B$_{PS}$ and PTP1B$_{PS*}$); LOV2 alters the IC$_{50}$ of the full-length PTP1B by 30%. For all figures, error bars denote SE for $n \geq 6$ independent reactions. Source data are provided as a Source Data file.

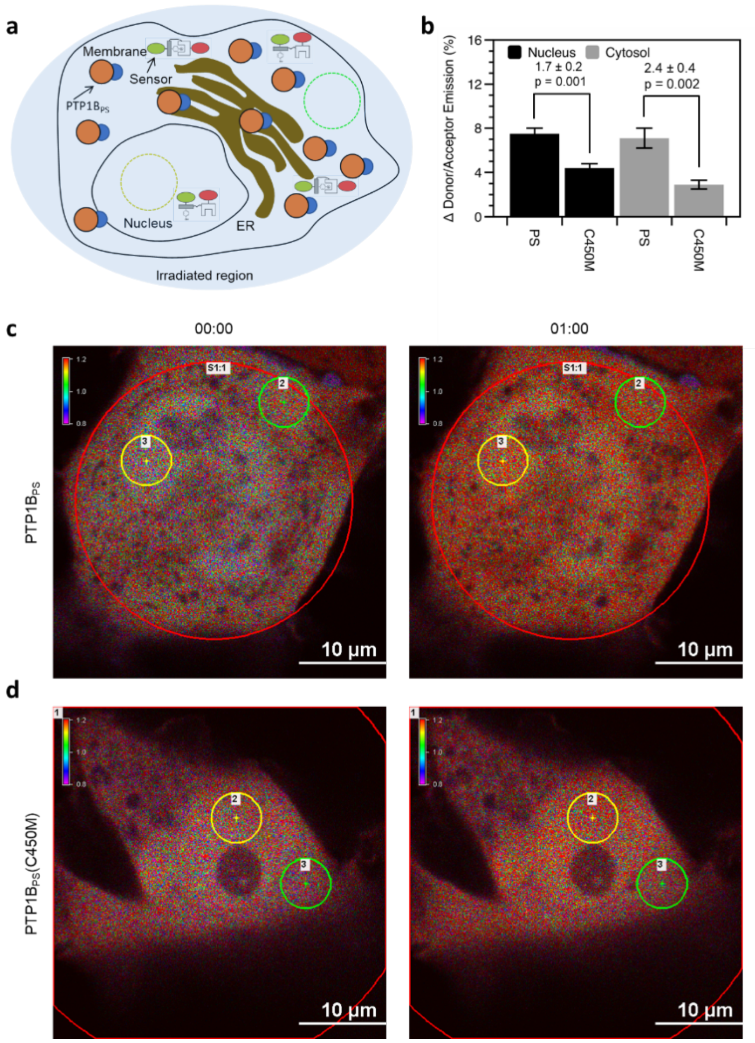


**Supplementary Figure 6 | Whole-cell irradiation of PTP1B$_{PS}$. a**, A COS-7 cell expressing both PTP1B$_{PS}$ and a biosensor is irradiated with blue light (457 nm). Circles depict the irradiated region (light blue oval), a nuclear region (yellow circle, d = 5 μm), and a cytosolic region (green circle, d = 5 μm). **b,** The percent change in donor/acceptor emission ratio over 1 min in nuclear and cytosolic regions. Each condition includes the interquartile average and SE for n = 11 biological replicates**. c**, Images of a COS-7 cell expressing both PTP1B$_{PS}$ and a biosensor at (left) 0 min after irradiation with blue light and (right) 1 min after irradiation. **d**, Images of a COS-7 cell expressing both PTP1B$_{PS}$(C450M) and a biosensor at (left) 0 min after irradiation with blue light and (right) 1 min after irradiation. Source data are provided as a Source Data file.

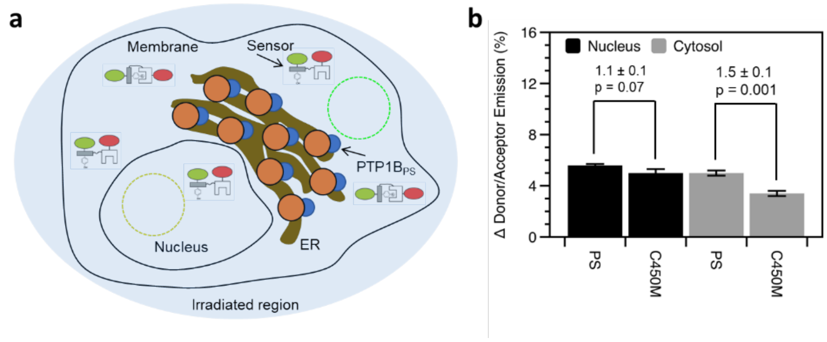


**Supplementary Figure 7 | Whole-cell irradiation of PTP1B$_{PS**}$. a,** A COS-7 cell expressing both PTP1B$_{PS**}$ and a biosensor is irradiated with blue light (457 nm). Circles depict the irradiated region (light blue oval), a nuclear region (yellow circle, d = 5 µm), and a cytosolic region (green circle, d = 5 µm). **b,** The percent change in donor/acceptor emission ratio measured over 1 min within the nuclear and cytosolic regions. Each condition includes the interquartile average and SE for n = 11 biological replicates. Source data are provided as a Source Data file.

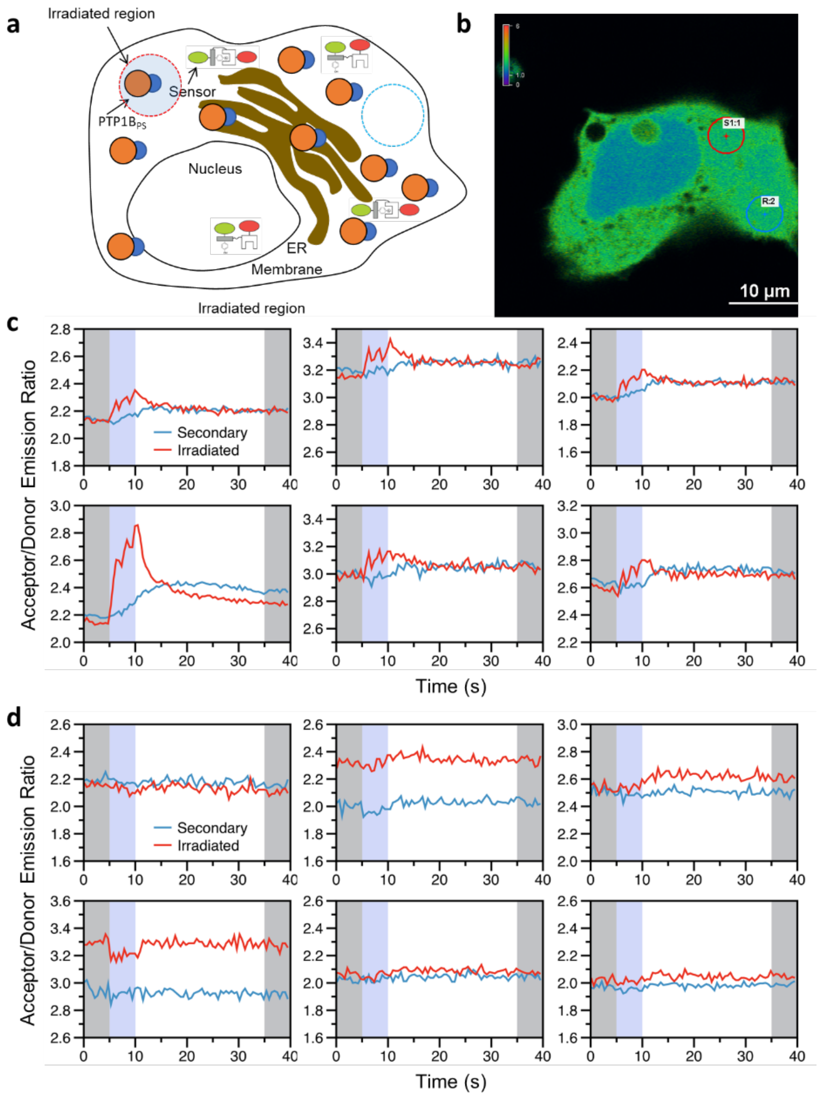


**Supplementary Figure 8 | Localized irradiation of PTP1B$_{PS}$. a**, A COS-7 cell expressing both PTP1B$_{PS}$ and a biosensor is locally irradiated with 405-nm light. Circles depict the irradiated region (red with blue interior, d = 5 µm) and a secondary region (blue, d = 5 µm) in the cytosol. **b**, Image of a COS-7 cell expressing both PTP1B$_{PS}$ and a biosensor; circles show irradiated and secondary regions. **c**, Time courses of donor/acceptor emission ratios in each region. **d**, Time courses of emission ratios measured in locally irradiated COS-7 cells expressing both PTP1B$_{PS}$(C450M) and the biosensor. In **c** and **d**, Shading highlights 5-s periods before (gray), during (blue), and after (gray) illumination. Source data are provided as a Source Data file.

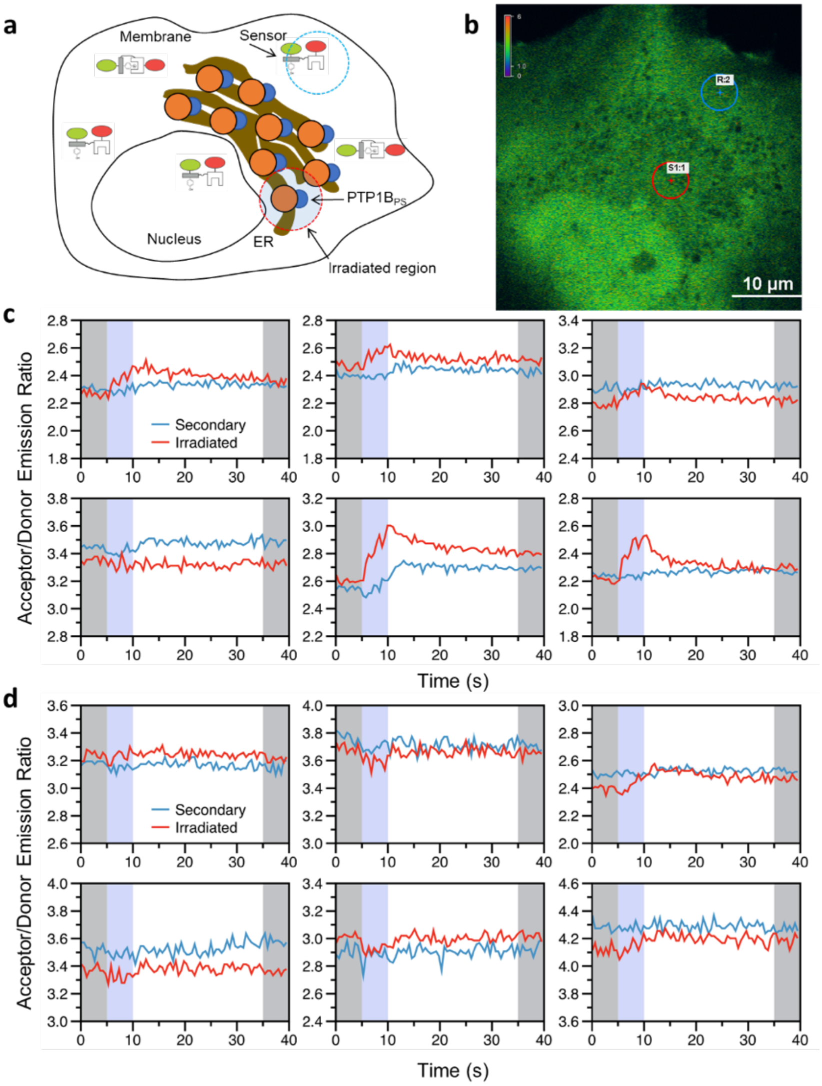


**Supplementary Figure 9 | Localized irradiation of PTP1B$_{PS**}$ near the nucleus. a**, A COS-7 cell expressing both PTP1B$_{PS**}$ and a biosensor is irradiated near the nucleus with 405-nm light. Circles depict the irradiated region (red with blue interior, d = 5 μm) and a secondary region (blue, d = 5 μm), both located within the cytosol. **b**, Image of a COS-7 cell expressing both PTP1B$_{PS**}$ and a biosensor; circles depict the irradiated and secondary regions. **c**, Time courses of donor/acceptor emission ratios measured in both regions **b**. **d**, Time courses of donor/acceptor emission ratios measured in locally irradiated COS-7 cells expressing both PTP1B$_{PS**}$(C450M) and the biosensor. In **c** and **d**, Shading highlights 5-s periods before (gray), during (blue), and after (gray) illumination. Source data are provided as a Source Data file.

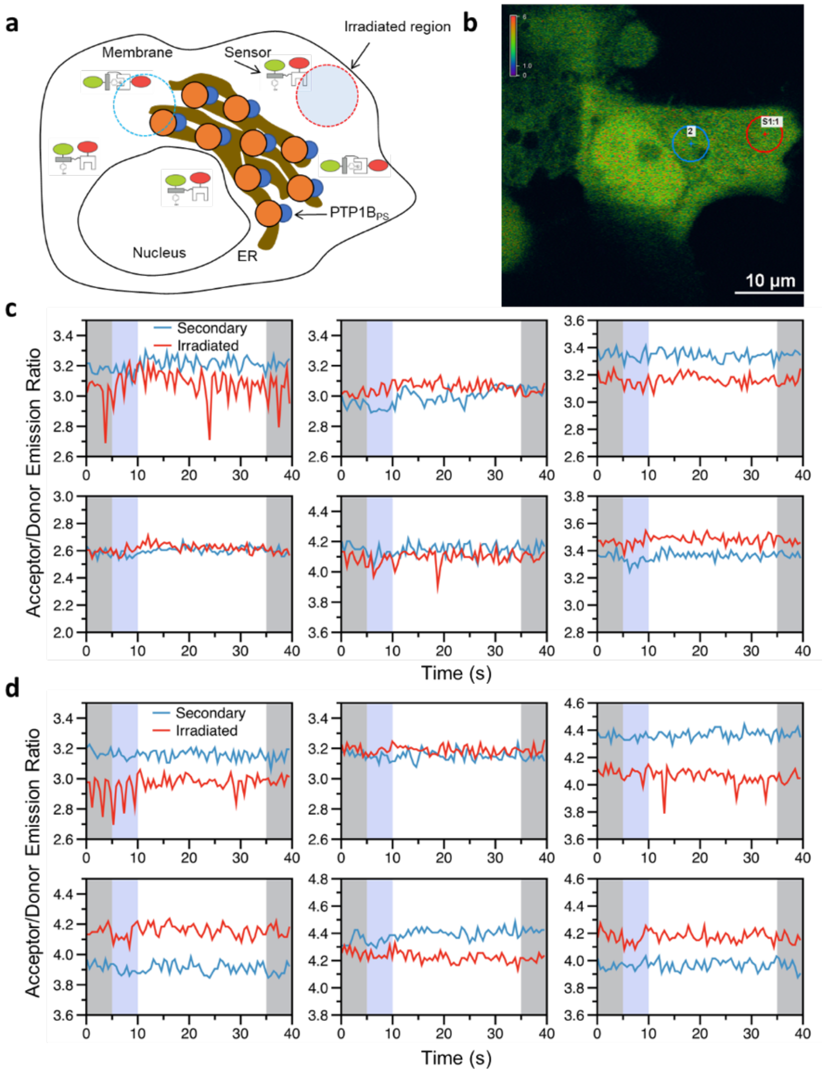


**Supplementary Figure 10 | Localized irradiation of PTP1B$_{PS**}$ near the plasma membrane.** **a**, A COS-7 cell expressing both PTP1B$_{PS**}$ and a biosensor is irradiated near the plasma membrane with 405-nm light. Circles depict the irradiated region (red with blue interior, d = 5 µm) and a secondary region (blue, d = 5 µm) in the cytosol. **b**, Image of a COS-7 cell expressing both PTP1B$_{PS**}$ and a biosensor. Circles ( d = 5 µm) show an irradiated region (red) and a secondary region (blue). **c**, Time courses of donor/acceptor emission ratios measured in both regions. **d**, Time courses of emission ratios measured in COS-7 cells expressing both PTP1B$_{PS**}$(C450M) and the biosensor. In **c** and **d**, Shading highlights 5-s periods before (gray), during (blue), and after (gray) illumination. Source data are provided as a Source Data file.

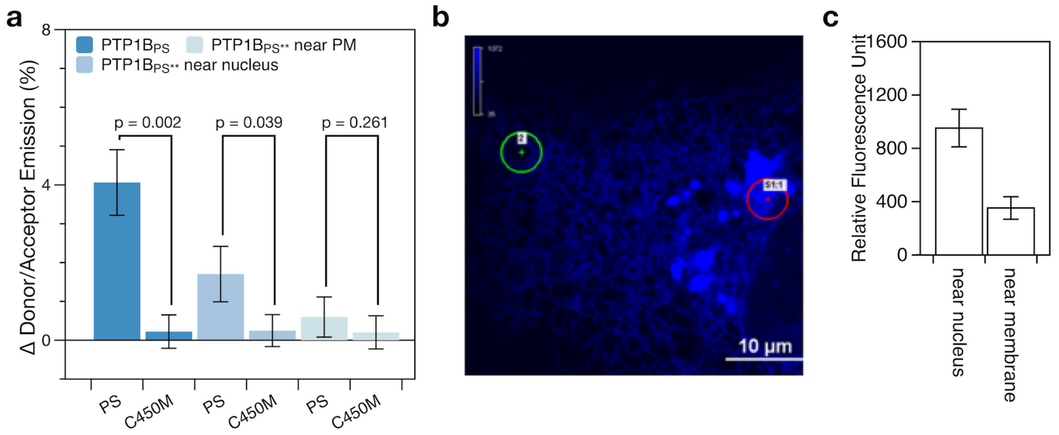


**Supplementary Figure 11 | Spatial heterogeneity of an ER-bound protein. a**, The change in FRET between 5-s intervals measured before and after illumination (gray regions in Figure 4e of the main text). For PTP1B$_{PS**}$, illumination near the nucleus, but not the plasma membrane (PM) causes a detectable change in FRET. **b**, Image of a COS-7 cell expressing an ER label[6]: BFP-Sec61β. Circular regions (d = 5 μm) appear near the nucleus (red) and plasma membrane (green). **c**, The average fluorescence of the nuclear region exceeds the average fluorescence of the membrane region by 2.7-fold, suggesting that the nuclear region contains more ER. Error bars denote SE for n = 6 (**a**) and n = 9 (**b**) biological replicates. Source data are provided as a Source Data file.

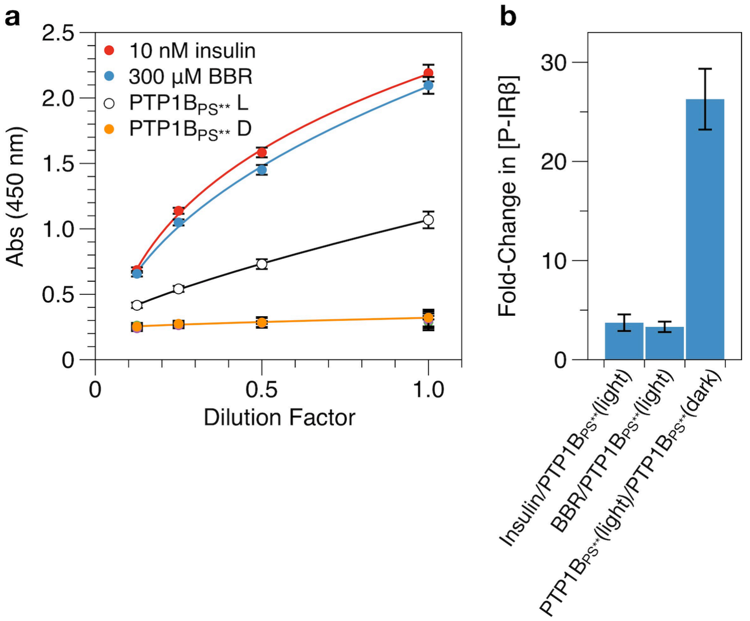


**Supplementary Figure 12 | Enzyme-linked immunosorbent assay (ELISA) of insulin receptor phosphorylation. a**, Measurements of insulin receptor phosphorylation (reported as a yellow color derived from 3,3',5,5'-tetramethylbenzidine) in cell lysate from either (i) HEK293T/17 cells exposed to 10 nM of insulin or 300 µM of BBR or (ii) PTP1B$_{PS**}$-expressing HEK293T/17 cells exposed to 455 nm light or kept in the dark. The dark dataset overlaps with those from (i) HEK293T/17 cells exposed to 1.5% DMSO (the concentration present in the BBR experiment) and (ii) PTP1B$_{PS**}$(C450M)-expressing HEK293T/17 cells exposed to 455 nm or kept in the dark. The overlap indicates that PTP1B$_{PS**}$ does not alter the native phosphorylation state of its regulatory targets. Curves denote fits to the four-parameter logistic equation: y = d+(a-d)/(1+(x/c)^b), where y is absorbance at 450 nm, and x is the sample dilution (Supplementary Table 9). Error bars denote SE with n = 3 biological replicates. **b**, Estimates of the difference in phosphorylated insulin receptor concentration between pairs of samples (Supplementary Table 10). Error bars represent SD of two independent estimates, each based on a separate standard curve from **a** (See Supplementary Note 2). Source data are provided as a Source Data file.

### Supplementary Table 1. Post-translational modifications of PTP1B.

| *Modification of PTP1B* | *Fold Change in Activity* | *Details of Measurement** | *Reference* |
|---|---|---|---|
| *Insulin-stimulated* ***phosphorylation*** *of tyrosines and dephosphorylation of serines* | 1.7-2.9 (inhibition) | Comparison of the activities of PTP1B immunoprecipitates collected from fat (2.9) and skeletal muscle (1.7) just before and 15 min after insulin stimulation of mice (Raytide-labeled substrate; p. 29523). | [7] |
| *Forskolin-stimulated* ***phosphorylation*** *of serines and dephosphorylation of tyrosines* | 1.7-1.8 (activation) | Comparison of the activities of immunoprecipitates collected from fat (1.8) and skeletal muscle (1.7) in the presence and absence of forskolin, an activator of adenylyl cyclase (p. 29524). | [7] |
| *Calpain-mediated* ***proteolysis*** *of disordered C-terminus* | 1.7 (activation) | Comparison of the activities of calpain-proteolyzed and wild-type PTP1B on p-nitrophenyl phosphate (p. 964). | [8] |
| *EGFR-mediated* ***phosphorylation*** *of Y66* | 3.1 (activation) | Comparison of the activities of EGFR-phosphorylated PTP1B and PTP1B(Y66F) on pNPP (p. 144). | [9] |
| *PIAS1-mediated* ***sumoylation*** *of K335 and K347 (primary sites)* | 2.3 (inhibition) | Comparison of the activities of PIAS1-sumoylated and wild-type PTP1B(1-403) on pNPP (p. 82). | [10] |

* Referenced figures and page numbers include descriptions of measurements and/or relevant source data; when necessary, we digitized the source data to obtain numerical values for DR by assuming a linear relationship between assay signal and enzyme activity.

**Supplementary Table 2. PTP1B-LOV2 Chimeras.**

| Control | WT | Note | Ref |
|---|---|---|---|
| ***Crossover*** | 1 | Analysis of crossover location | N/A |
| | 2 | Analysis of crossover location | |
| | 3 | Analysis of crossover location | |
| | 5 | Analysis of crossover location | |
| | 6 | Analysis of crossover location | |
| | 7 | Analysis of crossover location | |
| | 8 | Analysis of crossover location | |
| ***Partition*** | 7.1 | Shortening of SHEDLATTLE (α7A'α) to SHEDATTLE | |
| | 7.2 | Shortening of SHEDLATTLE (α7A'α) to SHEATTLE | |
| | 7.3 | Shortening of SHEDLATTLE (α7A'α) to SHATTLE | |
| | 7.4 | Shortening of SHEDLATTLE (α7A'α) to SATTLE | |
| | 7.5 | Shortening of SHEDLATTLE (α7A'α) to ATTLE | |
| | 7.6 | Shortening of SHEDLATTLE (α7A'α) to SHEDLTTLE | |
| | 7.7 | Shortening of SHEDLATTLE (α7A'α) to SHEDLTLE | |
| | 7.8 | Shortening of SHEDLATTLE (α7A'α) to SHEDLLE | |
| | 7.9 | Shortening of SHEDLATTLE (α7A'α) to SHEDLE | |
| | 7.10 | Shortening of SHEDLATTLE (α7A'α) to SHEDL | |
| ***Jα*** | 7.1(L514A) | Stabilization of Jα helix | |
| | 7.1(G528A) | Stabilization of Jα helix | |
| | 7.1(G528A/N538E) | Stabilization of Jα helix | |
| | 7.1ΔJα | Removal of Jα helix | |
| | 7.1(I532E) | Destabilization of Jα helix | |
| | 7.1(I539E) | Destabilization of Jα helix | |
| ***A'α*** | 7.1(T406A) | Stabilization of A'α helix | |
| | 7.1(T407A) | Stabilization of A'α helix | |
| | 7.1(T406A/T407A) | Stabilization of A'α helix | |
| | 7.1(L408D) | Destabilization of A'α helix | |
| ***α7*** | 7.1(H296A) | Stabilization of A'α helix | |
| | 7.1(E297A) | Stabilization of A'α helix | |
| | 7.1(D298A) | Stabilization of A'α helix | |
| | 7.1(S296A) | Stabilization of A'α helix | |
| | 7.1(D289A) | Stabilization of A'α helix | |
| | 7.1(K292A) | Stabilization of A'α helix | |
| | 7.1(E293A) | Stabilization of A'α helix | |
| ***Combination*** | 7.1(S286A/T406A) | Stabilization of α7A'α overlap | |
| | 7.1(K292A/T406A) | Stabilization of α7A'α overlap | |
| | 7.1(S296A/K292A) | Stabilization of α7A'α overlap | |
| | 7.1(S286A/K292A/T406A) | Stabilization of α7A'α overlap | |
| | 7.1(H296A/T406A) | Stabilization of α7A'α overlap | |
| ***Allostery*** | 7.1(Y152A/Y153A) | Destabilization of allosteric control | |
| ***Photoinactive mutant*** | 7.1(C450M) | Inactivation of photoswitching | |

### Supplementary Table 3. Primers used to assemble PTP1B-LOV2 chimeras.

| # | *PTP1B* | | *LOV2* | |
|---|---|---|---|---|
| | F Primer | R Primer | F Primer | R Primer |
| *1* | aaaaaccatggagatggaaaaggagttcgagc | ccgccttagctcatcatcaaaactctcgtcccccatgatgaatttggcac | gtgccaaattcatcatgggggacgagagttttgatgatgagctaaggcgg | tttttggatcctcagtggtggtggtggtggtgctcgagaagttcttttgccgcctcatcaat |
| *2* | aaaaaccatggagatggaaaaggagttcgagc | catttctttccgccttagctcatcatcaaaggaagagtcccccatgatgaatttgg | ccaaattcatcatgggggactcttcctttgatgatgagctaaggcggaaagaaatg | tttttggatcctcagtggtggtggtggtggtgctcgagaagttcttttgccgcctcatcaat |
| *3* | aaaaaccatggagatggaaaaggagttcgagc | cttctcatttctttccgccttagctcatcctgcacggaagagtccc | gggactcttccgtgcaggatgagctaaggcggaaagaaatgagaag | tttttggatcctcagtggtggtggtggtggtgctcgagaagttcttttgccgcctcatcaat |
| *5* | aaaaaccatggagatggaaaaggagttcgagc | gtagtagccaagtctataccccttctcatctccttccactgatcctgcacg | cgtgcaggatcagtggaaggagatgagaaggggtatagacttggctactac | tttttggatcctcagtggtggtggtggtggtgctcgagaagttcttttgccgcctcatcaat |
| *6* | aaaaaccatggagatggaaaaggagttcgagc | gtgtagtagccaagtctatacccccttctaagctccttccactgatcctgc | gcaggatcagtggaaggagcttagaaggggtatagacttggctactacac | tttttggatcctcagtggtggtggtggtggtgctcgagaagttcttttgccgcctcatcaat |
| *7* | aaaaaccatggagatggaaaaggagttcgagc | gttcttctcaatacgttcaagtgtagtagccaggtcctcgtgggaaagctc | gagctttcccacgaggacctggctactacacttgaacgtattgagaagaac | tttttggatcctcagtggtggtggtggtggtgctcgagaagttcttttgccgcctcatcaat |
| *8* | aaaaaccatggagatggaaaaggagttcgagc | gggtcagtaatgacaaagttcttctcaatacgctcgggtgggggctccag | ctggagcccccacccgagcgtattgagaagaactttgtcattactgaccc | tttttggatcctcagtggtggtggtggtggtgctcgagaagttcttttgccgcctcatcaat |
| *7.1* | aaaaaccatggagatggaaaaggagttcgagc | gttcttctcaatacgttcaagtgtagtagcgtcctcgtgggaaagctcc | gga gct ttc cca cga gga cgctactacacttgaacgtattgagaagaac | tttttggatcctcagtggtggtggtggtggtgctcgagaagttcttttgccgcctcatcaat |
| *7.2* | aaaaaccatggagatggaaaaggagttcgagc | gttcttctcaatacgttcaagtgtagtagcctcgtgggaaagctccttcc | ggaaggagctttcccacgaggctactacacttgaacgtattgagaagaac | tttttggatcctcagtggtggtggtggtggtgctcgagaagttcttttgccgcctcatcaat |
| *7.3* | aaaaaccatggagatggaaaaggagttcgagc | gttcttctcaatacgttcaagtgtagtagcgtgggaaagctccttccactg | cagtggaaggagctttcccacgctactacacttgaacgtattgagaagaac | tttttggatcctcagtggtggtggtggtggtgctcgagaagttcttttgccgcctcatcaat |
| *7.4* | aaaaaccatggagatggaaaaggagttcgagc | gttcttctcaatacgttcaagtgtagtagcggaaagctccttccactgatcc | ggatcagtggaaggagctttccgctactacacttgaacgtattgagaagaac | tttttggatcctcagtggtggtggtggtggtgctcgagaagttcttttgccgcctcatcaat |
| *7.5* | aaaaaccatggagatggaaaaggagttcgagc | gttcttctcaatacgttcaagtgtagtagcaagctccttccactgatcctgc | gcaggatcagtggaaggagcttgctactacacttgaacgtattgagaagaac | tttttggatcctcagtggtggtggtggtggtgctcgagaagttcttttgccgcctcatcaat |
| *7.6* | aaaaaccatggagatggaaaaggagttcgagc | caaagttcttctcaatacgttcaagtgtagtcaggtcctcgtgggaaagc | gctttcccacgaggacctgactacacttgaacgtattgagaagaactttg | tttttggatcctcagtggtggtggtggtggtgctcgagaagttcttttgccgcctcatcaat |

**Supplementary Table 3 (continued). Primers to assemble PTP1B-LOV2 chimeras.**

| # | ***PTP1B*** | | ***LOV2*** | |
|---|---|---|---|---|
| | F Primer | R Primer | F Primer | R Primer |
| *7.7* | aaaaaccatggagatggaaaaggagttcgagc | gacaaagttcttctcaatacgttcaagtgtcaggtcctcgtgggaaagc | gctttcccacgaggacctgacacttgaacgtattgagaagaactttgtc | tttttggatcctcagtggtggtggtggtggtgctcgagaagttcttttgccgcctcatcaat |
| *7.8* | aaaaaccatggagatggaaaaggagttcgagc | cagtaatgacaaagttcttctcaatacgttcaagcaggtcctcgtgggaaagc | gctttcccacgaggacctgcttgaacgtattgagaagaactttgtcattactg | tttttggatcctcagtggtggtggtggtggtgctcgagaagttcttttgccgcctcatcaat |
| *7.9* | aaaaaccatggagatggaaaaggagttcgagc | gtcagtaatgacaaagttcttctcaatacgttccaggtcctcgtgggaaagc | gctttcccacgaggacctggaacgtattgagaagaactttgtcattactgac | tttttggatcctcagtggtggtggtggtggtgctcgagaagttcttttgccgcctcatcaat |
| *7.10* | aaaaaccatggagatggaaaaggagttcgagc | ggtcagtaatgacaaagttcttctcaatacgcaggtcctcgtgggaaagc | gctttcccacgaggacctgcgtattgagaagaactttgtcattactgacc | tttttggatcctcagtggtggtggtggtggtgctcgagaagttcttttgccgcctcatcaat |
| *7.1ΔJα* | aaaaaccatggagatggaaaaggagttcgagc | gttcttctcaatacgttcaagtgtagtagcgtcctcgtgggaaagctcc | gga gct ttc cca cga gga cgctactacacttgaacgtattgagaagaac | tttttggatcctcagtggtggtggtggtggtgctcgaggacatgctcagttccatccaa |
| *$PTP1B_{281}$* | aaaaaccatggagatggaaaaggagttcgagc | tttttggatcctcaatgatgatgatgatgatgctcgaggatgaatttggcaccttcgatcac | N/A | N/A |

**Supplementary Table 4. Primers used for site-directed mutagenesis.**

| *Mutant* | *F Primer* | *R Primer* |
|---|---|---|
| *7.1(L514A)* | gtccagtactttattggggttcaggcggatggaactgagcatgtccgagat | atctcggacatgctcagttccatccgcctgaaccccaataaagtactggac |
| *7.1(G528A)* | gtccgagatgctgccgagagagaggcagtcatgctgattaagaaaactgca | tgcagttttcttaatcagcatgactgcctctctctcggcagcatctcggac |
| *7.1(G528A/ N538E)* | gtccgagatgctgccgagagagaggcagtcatgctgattaagaaaactgca/atgctgattaagaaaactgcagaagagattgatgaggcggcaaaagaactt | tgcagttttcttaatcagcatgactgcctctctctcggcagcatctcggac/aagttcttttgccgcctcatcaatctcttctgcagttttcttaatcagcat |
| *7.1(I532E)* | gccgagagagagggagtcatgctggagaagaaaactgcagaaaatattgat | atcaatattttctgcagttttcttctccagcatgactccctctctctcggc |
| *7.1(I539E)* | ctgattaagaaaactgcagaaaatgaggatgaggcggcaaaagaacttctc | gagaagttcttttgccgcctcatcctcattttctgcagttttcttaatcag |
| *7.1(T406A)* | agctttcccacgaggacgctgctacacttgaacgtattgagaa | ttctcaatacgttcaagtgtagcagcgtcctcgtgggaaagct |
| *7.1(T407A)* | tttcccacgaggacgctactgcacttgaacgtattgagaagaa | ttcttctcaatacgttcaagtgcagtagcgtcctcgtgggaaa |
| *7.1(T406A/ T407A)* | gagctttcccacgaggacgctgctgcacttgaacgtattgagaagaacttt | aaagttcttctcaatacgttcaagtgcagcagcgtcctcgtgggaaagctc |
| *7.1(L408D)* | ctttcccacgaggacgctactacagatgaacgtattgagaagaactttgtc | gacaaagttcttctcaatacgttcatctgtagtagcgtcctcgtgggaaag |
| *7.1(H296A)* | atcagtggaaggagctttccgcagaggacgctactacacttga | tcaagtgtagtagcgtcctctgcggaaagctccttccactgat |
| *7.1(E297A)* | agtggaaggagctttcccacgcagacgctactacacttgaacg | cgttcaagtgtagtagcgtctgcgtgggaaagctccttccact |
| *7.1(D298A)* | ggaaggagctttcccacgaggcagctactacacttgaacgtat | atacgttcaagtgtagtagctgcctcgtgggaaagctccttcc |
| *7.1(S286A)* | gccaaattcatcatgggggactctgccgtgcaggatcagtggaaggagctt | aagctccttccactgatcctgcacggcagagtcccccatgatgaatttggc |
| *7.1(D289A)* | atcatgggggactcttccgtgcaggctcagtggaaggagctttcccacgag | ctcgtgggaaagctccttccactgagcctgcacggaagagtcccccatgat |
| *7.1(K292A)* | gactcttccgtgcaggatcagtgggcggagctttcccacgaggacgctact | agtagcgtcctcgtgggaaagctccgcccactgatcctgcacggaagagtc |
| *7.1(E293A)* | tcttccgtgcaggatcagtggaaggcgctttcccacgaggacgctactaca | tgtagtagcgtcctcgtgggaaagcgccttccactgatcctgcacggaaga |
| *7.1(S286A/ T406A)* | gccaaattcatcatgggggactctgccgtgcaggatcagtggaaggagctt/agctttcccacgaggacgctgctacacttgaacgtattgagaa | aagctccttccactgatcctgcacggcagagtcccccatgatgaatttggc/ttctcaatacgttcaagtgtagcagcgtcctcgtgggaaagct |
| *7.1(K292A/ T406A)* | gactcttccgtgcaggatcagtgggcggagctttcccacgaggacgctact/gcggagctttcccacgaggacgctgctacacttgaacgtattgagaagaac | agtagcgtcctcgtgggaaagctccgcccactgatcctgcacggaagagtc/gttcttctcaatacgttcaagtgtagcagcgtcctcgtgggaaagctccgc |

**Supplementary Table 4 (continued). Primers used for site-directed mutagenesis.**

| *Mutant* | *F Primer* | *R Primer* |
|---|---|---|
| *7.1(S286A/K292A)* | gccaaattcatcatgggggactctgccgtgcaggatcagtggaaggagctt/gactctgccgtgcaggatcagtgggcggagctttcccacgaggacgctact | aagctccttccactgatcctgcacggcagagtcccccatgatgaatttggc/agtagcgtcctcgtgggaaagctccgcccactgatcctgcacggcagagtc |
| *7.1(S286A/K292A/T406A)* | gccaaattcatcatgggggactctgccgtgcaggatcagtggaaggagctt/gactctgccgtgcaggatcagtgggcggagctttcccacgaggacgctact/gcggagctttcccacgaggacgctgctacacttgaacgtattgagaagaac | aagctccttccactgatcctgcacggcagagtcccccatgatgaatttggc/agtagcgtcctcgtgggaaagctccgcccactgatcctgcacggcagagtc/gttcttctcaatacgttcaagtgtagcagcgtcctcgtgggaaagctccgc |
| *7.1(H296A/T406A)* | caggatcagtggaaggagctttccgccgaggacgctgctacacttgaacgt/agctttcccacgaggacgctgctacacttgaacgtattgagaa | acgttcaagtgtagcagcgtcctcggcggaaagctccttccactgatcctg/ttctcaatacgttcaagtgtagcagcgtcctcgtgggaaagct |
| *7.1(Y152A/Y153A)* | ttgatctctgaagatatcaagtcagcttatacagtgcgacagctagaattg | caattctagctgtcgcactgtataagctgacttgatatcttcagagatcaa |
| *7.1(C450M)* | cgtgaagaaattttgggaagaaacatgaggtttctacaaggtcctgaaact | agtttcaggaccttgtagaaacctcatgtttcttcccaaaatttcttcacg |
| *PTP1B$_{PS*}$ (L514A)* | gtccagtactttattggggttcaggcggatggaactgagcatgtccgagat | atctcggacatgctcagttccatccgcctgaaccccaataaagtactggac |
| *PTP1B$_{PS*}$ (G528A/N538E)* | gtccgagatgctgccgagagagaggcagtcatgctgattaagaaaactgca/atgctgattaagaaaactgcagaagagattgatgaggcggcaaaagaactt | tgcagttttcttaatcagcatgactgcctctctctcggcagcatctcggac/aagttcttttgccgcctcatcaatctcttctgcagttttcttaatcagcat |

### Supplementary Table 5. Primers used for variants of GFP-tagged $PTP1B_{PS}$.

| *Mutant* | *F Primer* | *R Primer* |
|---|---|---|
| *$PTP1B_{PS}$* | aaaaagaattcaatggagatggaaaaggagttcgagc | gaaaatattgatgaggcggcaaaagaacttggatccaaaaa |
| *$PTP1B_{PS**}$* | aaaaagaattcaatggagatggaaaaggagttcgagc | tttttggatccctatgtgttgctgttgaacaggaacctgtagcag aggtaagcgcc |
| *$PTP1B_{435}$* | aaaaagaattcaatggagatggaaaaggagttcgagc | tttttggatccctatgtgttgctgttgaacaggaacctgtagcag aggtaagcgcc |

### Supplementary Table 6. Primers used for Gibson assembly.

| # | *F Primer* | *R Primer* |
|---|---|---|
| *Pc#1-Biosensor* | ggctgctaacaaagcccgaaag | acatgatgatgatgatgatgcatggtatatc |
| *Pc#2-Biosensor* | catcatcatcatcatcatgtgagcaagg gcgaggag | tgatcttcccaaaataccacttgtacagctcgtccatg cc |
| *Pc#3-Biosensor* | tggtattttgggaagatcactcgtc | gagctccccctgtaggtgg |
| *Pc#4-Biosensor* | tacagggggagctcgtgtctaagggcg aagagctg | gctttgttagcagccttacttgtacagctcgtccatccc |
| *Pc#1-$PTP1B_{PS*}$* | gcggcaaaagaacttctggagcccccca cccg | atgatgatgatgctcgaggtaactcagtgcatggtcc tcgtc |
| *Pc#2-$PTP1B_{PS*}$* | ctcgagcatcatcatcatcatcattgagg | aagttcttttgccgcctcatcaat |
| *Pc#3-$PTP1B_{PS*}$* | tccagtgatcgaagttaggctgg | ctaacttcgatcactggaccgctg |
| *Pc#1-Biosensor-P2A-$PTP1B_{PS}$* | ggtggcgaccggtagcg | ctgatcataatcagccataccacatttgtagag |
| *Pc#2-Biosensor-P2A-$PTP1B_{PS}$* | ggcagcggcgccacc | gcttttctgctcccacaccatctc |
| *Pc#3- Biosensor-P2A-$PTP1B_{PS}$* | tgggagcagaaaagcaggggtgtcgt catgctcaac | gtatggctgattatgatcagttaaagttcttttgccgcc tcatcaatattttc |
| *Pc#4- Biosensor-P2A-$PTP1B_{PS}$* | ccggtcgccaccatggtgagcaaggg cgaggagc | cgccgctgcccttgtacagctcgtccatcccac |
| *Pc#4- Biosensor-P2A-$PTP1B_{PS**}$* | ccggtcgccaccatggtgagcaaggg cgaggagc | atggctgattatgatcagttatgtgttgctgttgaacag gaacc |

Note: Pc#1 denotes the first piece of a Gibson assembly.

### Supplementary Table 7. DNA fragments used for Gibson assembly

| # | *DNA sequence* |
|---|---|
| *P2A-PTP1B* | ggcagcggcgccaccaacttctccctgctgaagcaggccggcgacgtggaggagaaccccggc<br>cccatggagatggaaaaggagttcgagcagatcgacaagtccgggagctgggcggccatttacca<br>ggatatccgacatgaagccagtgacttcccatgtagagtggccaagcttcctaagaacaaaaaccga<br>aataggtacagagacgtcagtccctttgaccatagtcggattaaactacatcaagaagataatgactat<br>atcaacgctagtttgataaaaatggaagaagcccaaaggagttacattcttacccagggccctttgcct<br>aacacatgcggtcacttttgggagatggtgtgggagcagaaaagc |

### Supplementary Table 8. Optogenetic Constructs

| Protein* | Light-sensitive Moiety | Mechanism | Dynamic Range (DR) | Assay for DR** . | Cellular Response** | Ref |
|---|---|---|---|---|---|---|
| Pyruvate kinase M2 isoform | LOV2 from *A. sativa* phototropin I | Optically induced conformational change in LOV2 causes an activity-distorting conformational change in an enzyme. | 1.4-1.9 (activation) | Pyruvate production (in the presence and absence of light) measured by time-resolved series of 1H-NMR spectra (p. 2966). | In the presence of [U C-13] glucose, irradiation of HeLa cells increased C-13 incorporation into pyruvate (52% in light vs. 35% in dark; p. 2966). | 11 |
| RhoA GTPase | LOV2 from *A. sativa* phototropin I | Optically induced conformational change in LOV2 causes an activity-distorting conformational change in an enzyme. | 1.7 (inhibition) | Biosensor-based quantification (i.e., FRET) of GTPase activity in HeLa cells for dark- and light-state mutants of LOV2 (p. 1442). | Irradiation of fibroblasts showed increased protrusions upon irradiation (Supplementary Fig. 13). | 12 |
| Peptide inhibitor of protein kinase A (PKA) | LOV2 from *A. sativa* phototropin I | Optically induced unwinding of the Jα exposes an attached inhibitory peptide. | 1.9 (inhibition / binding) | Quantification of a Western blot showing the phosphorylation state of a PKA substrate before/after illumination (p. 790). | Irradiation of cells caused 2-fold reduction in forskolin-dependent phosphorylation of ERB (p. 790). | 13 |
| Vav2 GTP exchange factor | LOV2 from *A. sativa* phototropin I | Optically induced conformational change in LOV2 causes an activity-distorting conformational change in an enzyme. | 2.1 (inhibition) | Biosensor-based quantification (i.e., FRET) of Rac1 GTPase activity in HeLa cells for dark- and light-state mutants of LOV2 (p. 1443). | Irradiation of HeLa cells produced rapid reversible retraction (p. 1443). | 12 |
| Nuclear export sequence | LOV2 from *A. sativa* phototropin I | Optically induced unwinding of an internal Jα-A'α connection within a LOV2-LOV2 fusion exposes an embedded nuclear export sequence that can bind to CRM1. | 2.3 (binding) | Quantification of optically induced relocalization of a fluorescent protein from the nucleus to the cytosol (p. 399). | N/A | 14. |
| Peptide binding partner of engineered PDZ domain. | LOV2 from *A. sativa* phototropin I | Optically induced unwinding of the Jα helix uncages a peptide epitope and enables binding to a cognate PDZ domain. | 2.7 (binding) | Quantification of optically induced relocalization from the cytosol to the plasma membrane (Supplementary Fig. 6). | Irradiation of HeLa cells enabled localization of ePDZb1-mCherry to the plasma membrane and outer membrane of the mitochondria (p. 380). | 15 |

| Protein* | Light-sensitive Moiety | Mechanism | Dynamic Range (DR) | Assay for DR** . | Cellular Response** | Ref |
|---|---|---|---|---|---|---|
| Src kinase | LOV2 from *A. sativa* phototropin I | Optically induced conformational change in LOV2 causes an activity-distorting conformational change in an enzyme. | 3.0-5.2 (inhibition) | Quantification of (i) paxillin phosphorylation in *in vitro* assay (5.2) and (ii) tyrosine phosphorylation in cell lysate (3.0) before/after illumination (p. 1442). | Irradiation of SYF cells caused Src to translocate to focal adhesions; edge movements tended to increase, and cells polarized (p. 1442). | 12 |
| Peptide inhibitor of myosin light chain kinase | LOV2 from *A. sativa* phototropin I | Optically induced unwinding of the Jα exposes an attached inhibitory peptide. | 3.3 (inhibition / binding) | Quantification of a Western blot showing the phosphorylation state of myosin light chain before/after illumination (p. 791). | Irradiation of cells caused a statistically significant reduction in membrane protrusion velocity (p. 791). | 13 |
| Cdc42 GTPase | LOV2 from *A. sativa* phototropin I | Optically induced conformational change in LOV2 causes an activity-distorting conformational change in an enzyme. | 3.5 (inhibition) | Biosensor-based quantification (i.e., FRET) of GTPase activity in HeLa cells for dark- and light-state mutants of LOV2 (p. 1442). | Irradiation of fibroblasts produced retraction and an accumulation of actin at the cell periphery (Supplementary Fig. 13). | 12 |
| BphP1 from *Rhodopseudomonas palustrus* | BphP1 from *Rhodopseudomonas palustrus* | Optically induced conformational change in BphP1 changes its affinity for its binding partner PpsR2 | 3.5 (binding) | Binding: Quantification of optically induced relocalization of a fluorescent protein from the cytosol to the nucleus (p. 594). | Irradiation of Hela cells enabled 40-fold higher expression of SEAP under 740 nm light than in the dark (p. 594). Note: Transcription and translation are amplification reactions. | 16 |
| Nuclear export sequence | LOV2 from *A. sativa* phototropin I | Optically induced unwinding of the Jα exposes an embedded nuclear export sequence that can bind to CRM1. | 4 - 14 (binding) | Quantification of optically induced relocalization of a fluorescent protein from the nucleus to the cytosol (p. 399). The dynamic range differed between NESs. | Illumination of yeast cells expressing Bre1 bound to the photoswitchable construct caused a measurable decrease in H2B histone ubiquitylation (p. 400). | 14 |
| Rac1 GTPase | LOV2 from *A. sativa* phototropin I | Optically induced conformational change in LOV2 causes an activity-distorting conformational change in an enzyme. | 4.5 (inhibition) | Biosensor-based quantification (i.e., FRET) of GTPase activity in HeLa cells for dark- and light-state mutants of LOV2 (p. 1442). | Irradiation of fibroblasts caused reversible cell edge retraction and broad retraction of entire lamellae (p. 1443). | 12 |

| **Protein*** | **Light-sensitive Moiety** | **Mechanism** | **Dynamic Range (DR)** | **Assay for DR**** . | **Cellular Response**** | **Ref** |
|---|---|---|---|---|---|---|
| Fibroblast growth factor receptor 1 (FGFR1) | LOV from *V. frigida* aureochrome I | Optically induced dimerization of LOV2 induces dimerization--and activation--of the RTKs to which they are attached. | 5.5 (activation / transcription) | Quantification of MAPK/ERK pathway activation and subsequent luciferase expression (reporter) in HEK293 cells in response to blue light (p. 1716). | Irradiation of M38K cells induced phosphorylation of the receptor and ERK1/2 and caused an increase in cell proliferation and induced EMT-like morphological and gene expression changes (p. 1718). | 17 |
| *trp* repressor from *E. coli* | LOV2 from *A. sativa* phototropin I | Optically induced unwinding of the Jα helix causes TrpR domains to adopt conformations that enable binding to DNA. | 5.6 (binding) | Binding affinity measured through the quantification of RsaI protection by Trp in the presence and absence of light (p. 10712). | N/A | 18. |
| Adenylate cyclase (AC) from *Nostoc* sp. | BphG1 from *Rhodobacter sphaeroides* | Optically induced dimerization of BphG1 induces dimerization of AC, which functions as a homodimer. | 5.7 (activation / binding) | *In vitro* measurement of the kinetics of cAMP accumulation in the presence and absence of light (p. 10170) | Cholinergic neurons expressing the photoswitchable AC exhibited a higher frequency of body bends in the presence of red light (relative to the dark, p. 10170). | 19 |
| Cas9 endonuclease | LOV2 domain from *Rhodobacter sphaeroides* | Optically induced dissociation of a LOV2 dimer uncages a variant of Cas9 with LOV2 embedded within it, enabling endonuclease activity. | 6.5 (activation, transcription) | Quantification of dCas9-mediated transcriptional repression of RFP gene (p. 10006). | Irradiation of *E. coli* harboring photoswitchable cas9 yielded 3-fold repression of endogenous *lacZ* (p. 10011). | 20 |
| Raf1 kinase | Dronpa from *Pectinidae* coral | Optically induced dissociation of a Dronpa dimer uncages the active site. | 6.8 (activation) | Quantification of a Western blot showing the light-dependent phosphorylation ERK (p. 2). | A pulse of 500-nm light elicited transient increase in MEK phosphorylation, evidencing negative feedback on Raf activity (p. 4). Exposure of *C. elegans* to 500-nm light led to tail swelling with 73-79% penetrance (p. 5). | 21 |

| Protein* | Light-sensitive Moiety | Mechanism | Dynamic Range (DR) | Assay for DR** . | Cellular Response** | Ref |
|---|---|---|---|---|---|---|
| AcrIIA4 from *l. monocytogenes* prophage | LOV2 from *A. sativa* phototropin I | Optically induced conformational change in LOV2 causes an activity-distorting conformational change in AcrIIA4, reducing its ability to inhibit Cas9. | 7.4 (unbinding, transcription) | Quantification of dCas9-mediated transcriptional repression of luciferase gene in HEK293T cells (p. 925). | Irradiation of HEK293T cells yielded light-dependent insertion/deletion (indel) mutations (p. 925) and light-dependent telomere recruitment (p. 927) | 22 |
| SsrA peptide | LOV2 from *A. sativa* phototropin I | Optically induced unwinding of the Jα exposes an embedded inhibitory peptide. | 8 (binding) | Dark- and light-state binding affinity measured with a fluorescence polarization competition assay (p. 511). | N/A | 23 |
| MEK2 kinase | Dronpa from *Pectinidae* coral | Optically induced disassociation of a Dronpa dimer uncages the active site. | 8.6 (activation) | Quantification of a Western blot showing the light-dependent phosphorylation ERK (p. 2). | N/A | 21 |
| MEK1 kinase | Dronpa from *Pectinidae* coral | Optically induced dissociation of a Dronpa dimer uncages the active site. | 9.1 (activation) | Quantification of a Western blot showing the light-dependent phosphorylation ERK (p. 2). | Irradiation of cells with 500-nm light induced translocation pf ERK KTR-mRuby2 from nucleus to cytosol; irradiation with 400 nm light reversed this effect (p. 2). Exposure of C. elegans to 500-nm light produced tail swelling with 73-79% penetrance (p. 5). | 21 |
| Rac1 GTPase | LOV2 from *A. sativa* phototropin I | Optically induced unwinding of the Jα exposes active site of GTPase. | 10 (activation) | Binding affinity for PAK measured with isothermal titration calorimetry on light- and dark-state mutants (p. 104). | Irradiation of HeLa cells produced lamellipodial protrusions and membrane ruffles (p. 104). Irradiation of MEF cells at a spot near the cell edge produces local protrusion and distal retraction (p. 104). | 1 |
| Cre Recombinase | Cryptochrome 2 from *A. thaliana* | Optically induced dimerization of Cry2 and CIB permits assembly of a split recombinase. | 13.7 (activation) | Quantification of Cre recombinase activity in HEK293 cells pulsed with light (p. 428). | Brief irradiation of HEK293 cells enabled a ~14% increase in recombination (relative to the dark; p. 428). | 24 |

| Protein* | Light-sensitive Moiety | Mechanism | Dynamic Range (DR) | Assay for DR** . | Cellular Response** | Ref |
|---|---|---|---|---|---|---|
| Peptide binding partner (SsrA) of SspB. | LOV2 from *A. sativa* phototropin I | Optically induced unwinding of the Jα helix uncages the SsrA peptide, allowing it to bind to SspB. | 36 (binding) | Light- and dark-state binding affinity for SspB measured with a fluorescence polarization competition assay (p. 114). | Irradiation of fibroblasts expressing mitochondria-anchored LOV2-SsrA caused recruitment of SspB (page 115). Irradiation of fibroblasts expressing (i) membrane-anchored LOV2-SsrA and (i) a fusion of the DH/PH domain of Tiam1 to the N-terminus of TagRFP-SspB produced ruffles and lamellipodia formation (p. 116). | 25 |
| Vinculin-binding peptide from ipaA | LOV2 from *A. sativa* phototropin I | Optically induced unwinding of the Jα exposes an embedded inhibitory peptide. | 49 (binding) | Dark- and light-state binding affinity measured with a fluorescence polarization competition assay (p. 511). | Irradiation of *S. cerevisiae* expressing (i) LOV-ipaA linked to the Gal4 activation domain and (ii) vinculin D1 to the Gal4 binding domain enabled 12-fold higher expression of LacZ (relative to dark; p. 513). | 23 |
| LOV2 from *A. sativa* phototropin I | LOV2 from *A. sativa* phototropin I | Optically induced unwinding of the Jα helix prevents binding to Zdk, a protein evolved to bind the dark-state of LOV2. | > 150 (binding) | Binding affinity of dark- and light-state mutants measured with radiometric binding assay (p. 755). | Irradiation of HeLa cells expressing both (i) Zdk bound to constitutively active mutants of RhoA, Rac1, and Vav2 and (ii) mitochondria-localized LOV2 caused changes in edge velocity, ruffling, cell area, and protrusion distribution (p. 757). | 26 |

†This table provides a reference set of optogenetic constructs, each chosen for its reliance on a LOV domain or for its demonstrated use in signaling studies.
*Constructs in which the light-sensitive moiety exerts direct control over enzyme activity appear in blue; constructs in which the moiety controls protein-protein interactions appear in orange.
** Referenced figures and page numbers include descriptions of measurements and/or relevant source data; when necessary, we digitized the source data to obtain numerical values for DR.

### Supplementary Table 9. Four-parameter logistic regression

| *Parameter** | *PTP1B$_{PS**}$ (light)* | *Insulin (10 nM)* | *BBR (300 μM)* | *PTP1B$_{PS**}$ (dark)* |
|---|---|---|---|---|
| ***a*** | 0.247 | -1.427 | -0.370 | 0.234 |
| ***b*** | 0.753 | 0.279 | 0.419 | 0.675 |
| ***c*** | 10100 | 1840 | 11600 | 316000 |
| ***d*** | 850 | 31.7 | 127 | 445 |

[*]Fits of the dilution data displayed in Supplementary Fig. 12 to the four-parameter logistic equation: y = d+(a-d)/(1+(x/c)^b), where y is absorbance at 450 nm, and x is the fractional sample dilution (with 1 corresponding to no dilution). Fits to each of the four sample sets appear as lines in the figure.

### Supplementary Table 10. Differences in insulin receptor phosphorylation

| *Parameter** | *PTP1B$_{PS**}$ (light) Curve* | *Reference Curve* | *Average* | *SD* |
|---|---|---|---|---|
| *PTP1B$_{PS**}$ (light)/ PTP1B$_{PS**}$ (light)* | 1 | N/A | N/A | N/A |
| *PTP1B$_{PS**}$ (light)/ PTP1B$_{PS**}$ (dark)* | 28.4 | 24.1 | 26.3 | 3.1 |
| *Insulin (10 nM)/ PTP1B$_{PS**}$ (light)* | 4.3 | 3.1 | 3.3 | 0.5 |
| *BBR (300 μM)/ PTP1B$_{PS**}$ (light)* | 3.7 | 2.9 | 3.7 | 0.8 |

[*]We used the fits described in Supplementary Table 9 to estimate the fold-difference in concentration of phosphorylated insulin receptor between two samples: (i) PTP1B$_{PS**}$-expressing HEK293T/17 cells exposed to 455 nm light and (ii) a second sample. In the first column, we based these estimates on the dilution curve for the first sample; in the second column, we based them on the dilution curve of the second sample. Consistency between the two estimates suggests that they provide good quantitative metrics (See Supplementary Note 2).
